## Supplemental tumor model for "PhysiCell Studio: a graphical tool to make agent-based modeling more accessible"

### Supplemental material.


Supplemental material.
- Add pressure mechanofeedback on cycling
- Add oxygen-based cycling
- Add hypoxia-driven necrosis
- Add a cytotoxic drug
- Add release of dead cell debris
- Add macrophages
- Add pro-inflammatory factor
- Add effector T cells
- Intracellular modeling


We continue with the demonstration on how one might use PhysiCell Studio by adding more details to the tumor model. For more information on why certain parameter values were chosen, refer to Part 2 slides at github.com/physicell-training/nw2023.

### Add pressure mechanofeedback on cycling

We use the Rules tab to dynamically modify the cell cycle (behavior) as a function of cell pressure (signal). Specifically, we want an increase in pressure to decrease the transition rate of the ("live") cell cycle. The modeling grammar[1] used in the Rules tab was introduced in PhysiCell version 1.12.0.

In the "Rules" tab:

- the top "Cell Type" combobox should only contain "cancer"
- choose "pressure" as the Signal
- choose "cycle entry" as the Behavior
- choose "decreases" next to Behavior
- enter 0 as the Saturation value of Behavior
- enter 4 as the Hill power
- enter 1 as the Half-max
- optionally, you can click "Plot" (next to "Add rule") to see the Hill function (Figure 1)
- click "Add rule" (rule should appear in the table at row #1, Figure 2)
- at the bottom of the tab, set rules file name to "cell_rules.csv"
- be sure "enable" is checked

- optionally, click “Save” button to save the rule to “cell_rules.csv” (otherwise, it will be automatically saved when you Run a simulation)
- in the Run tab, “Run simulation”

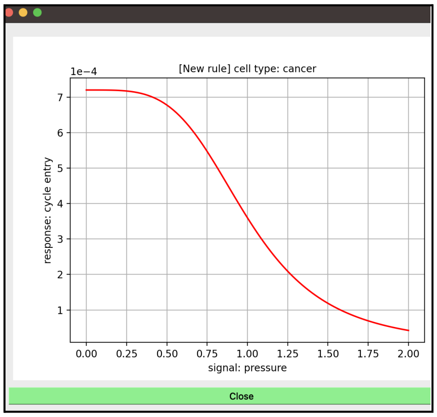


Figure 1. (pressure, cycle entry) Hill function.

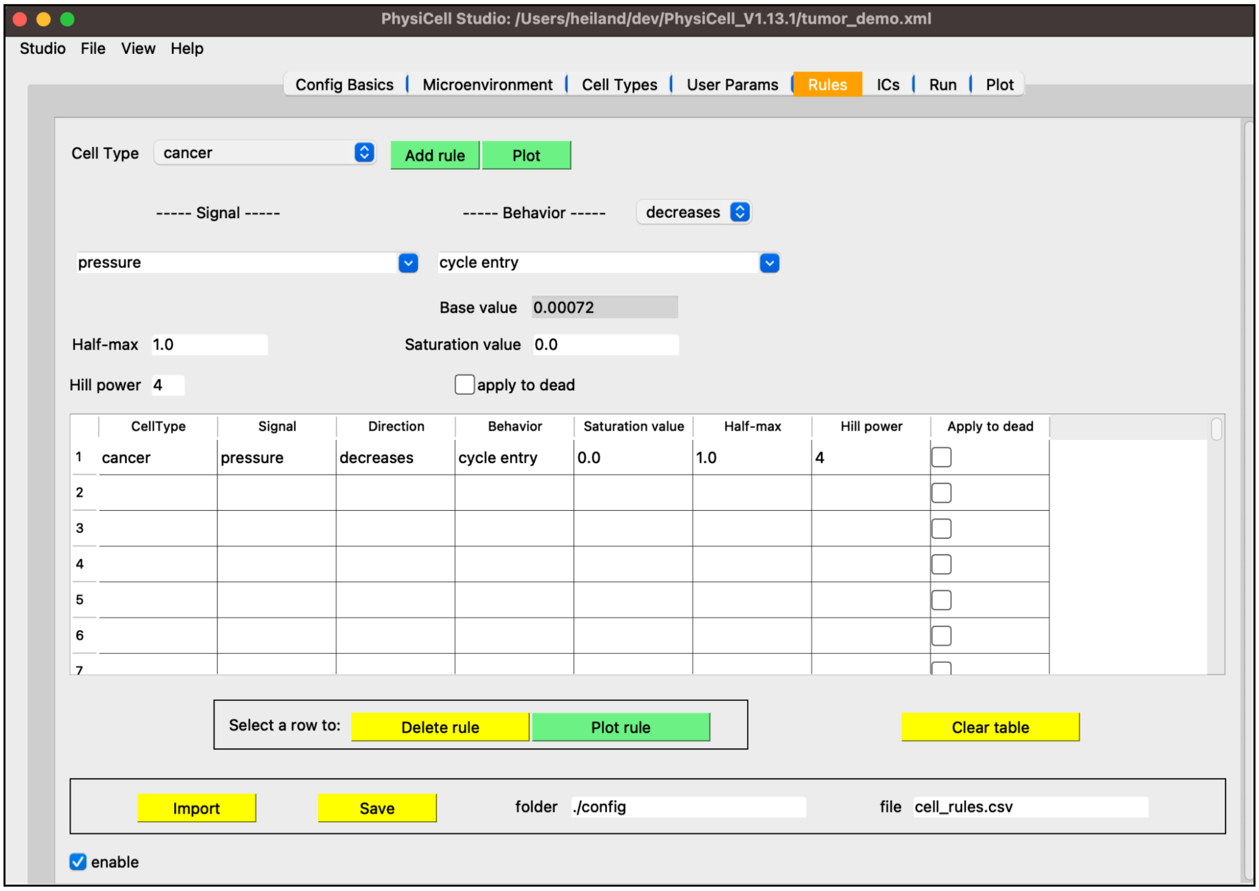


Figure 2. Rule #1: Pressure decreases cycle entry.

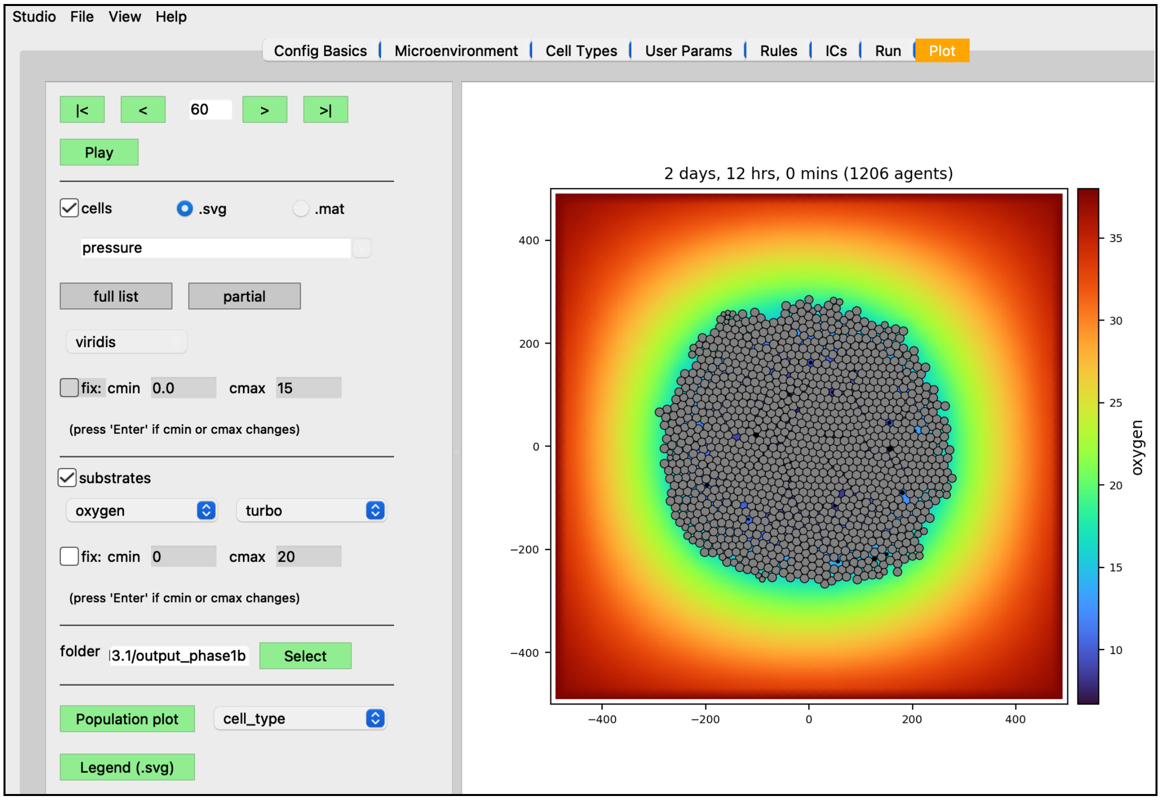


Figure 3. Results with one rule.

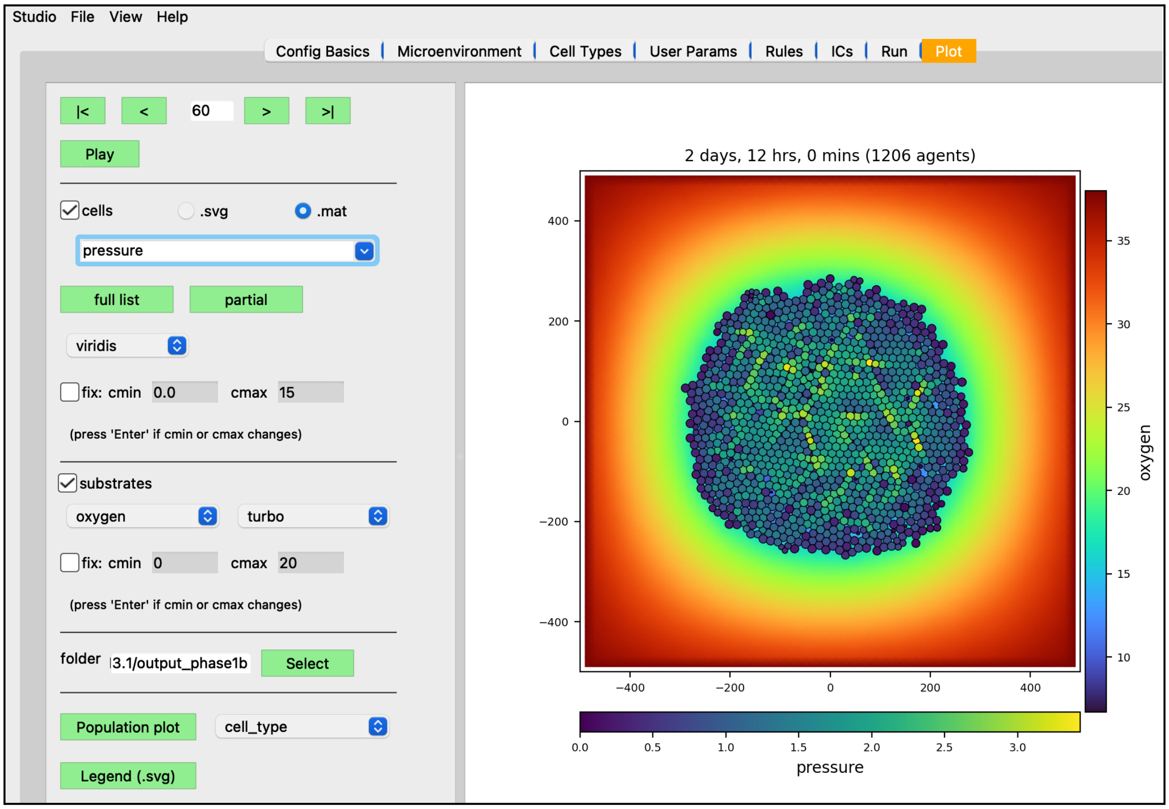


Figure 4. Coloring cells with pressure values

### Add oxygen-based cycling

- in the “Cell Types” tab, “Cycle” sub-tab
    - select “transition rate(s)” radio button
    - enter 0 for the rate

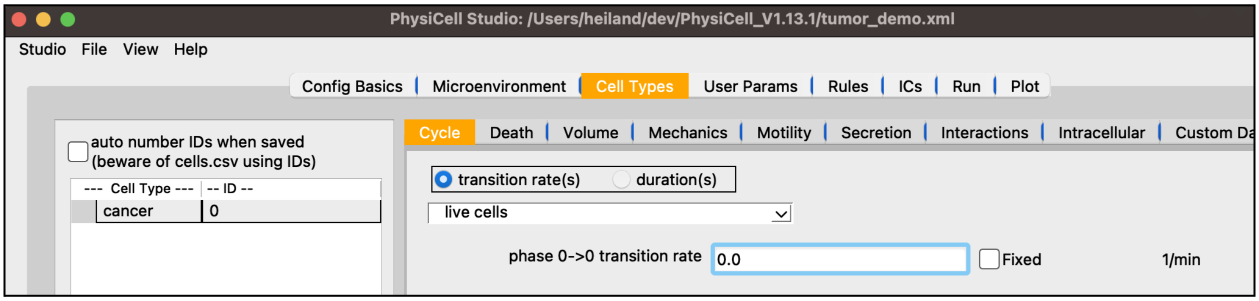


Figure 5. Set Cycle transition rate to 0.

- in the “Rules” tab, the “Cell Type” combobox should only contain “cancer”
- choose “oxygen” as the Signal
- chose “cycle entry” as the Behavior
- choose “increases” as the response
- choose 0.00093 as the Saturation value of the Behavior
- choose 21.5 (mmHg) as the Half-max
- choose 4 as the Hill power
- optionally, Plot the new rule
- click “Add rule” and “Save”
- Run a new simulation

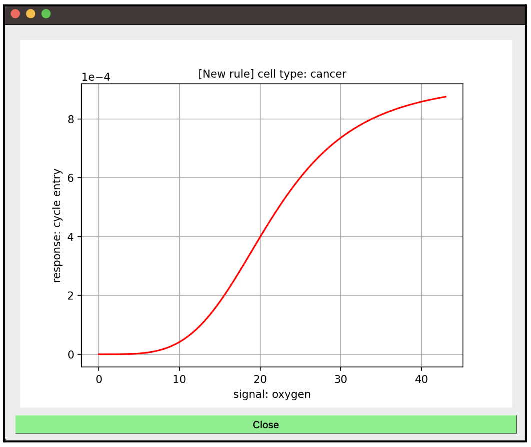


Figure 6. (oxygen, cycle entry) Hill function.

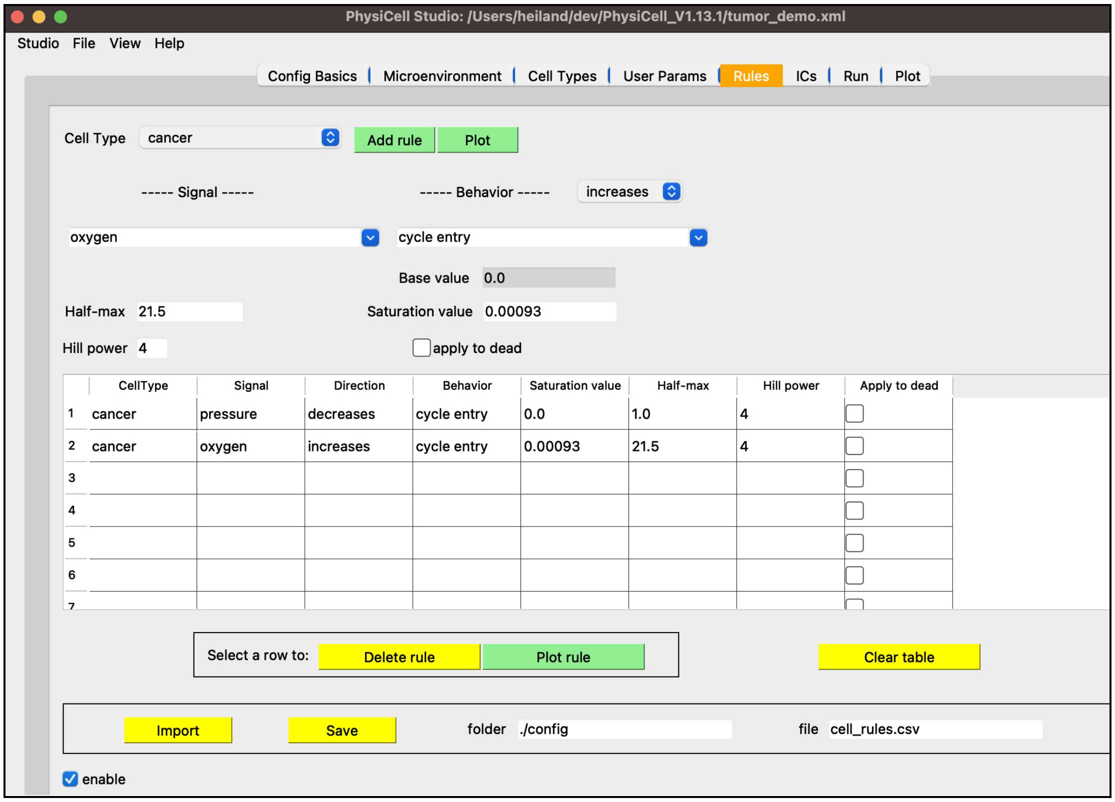


Figure 7. Rule #2: Oxygen increases cycle entry

After running a new simulation with this second rule, we visualize results as follows:

- in the "Plot" tab, select the ".mat" radio button for cells
- click "full list" to show all options for cell scalar values
- select "current_cycle_phase_exit_rate" as the scalar value
- select the "jet" colormap for cells
- check "fix" to limit the colormap ranges, with cmin=0 and cmax=0.00093 (press Enter)
- select the "YlOrRd" colormap for the substrate
- click "Play" to show the simulation
- notice the greatest cycling is on the outer periphery of the tumor

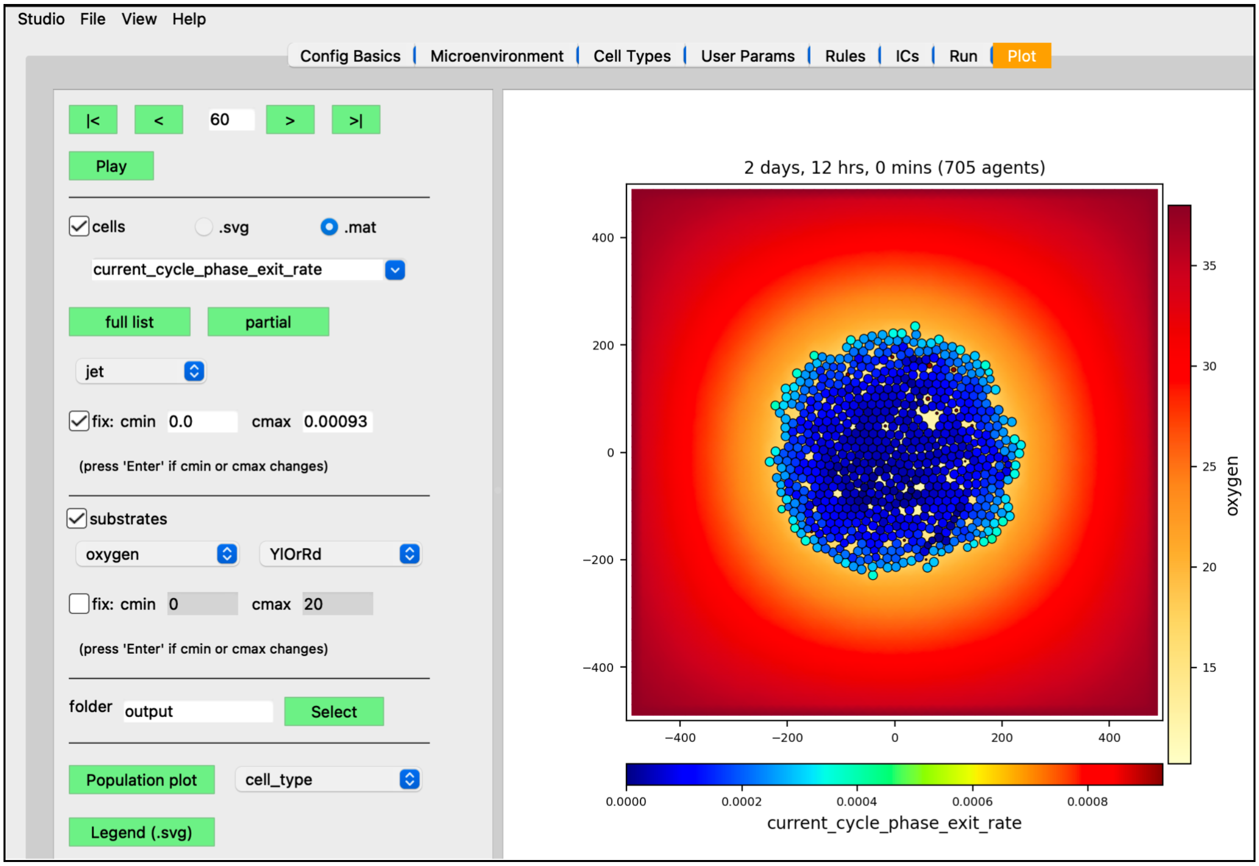

Figure 8. Results with two rules (coloring cells with .mat cycle phase exit rate)

### Add hypoxia-driven necrosis

- in the “Cell Types” tab, “Death” sub-tab, “Necrosis” section
  - set “death rate” to 3e-3

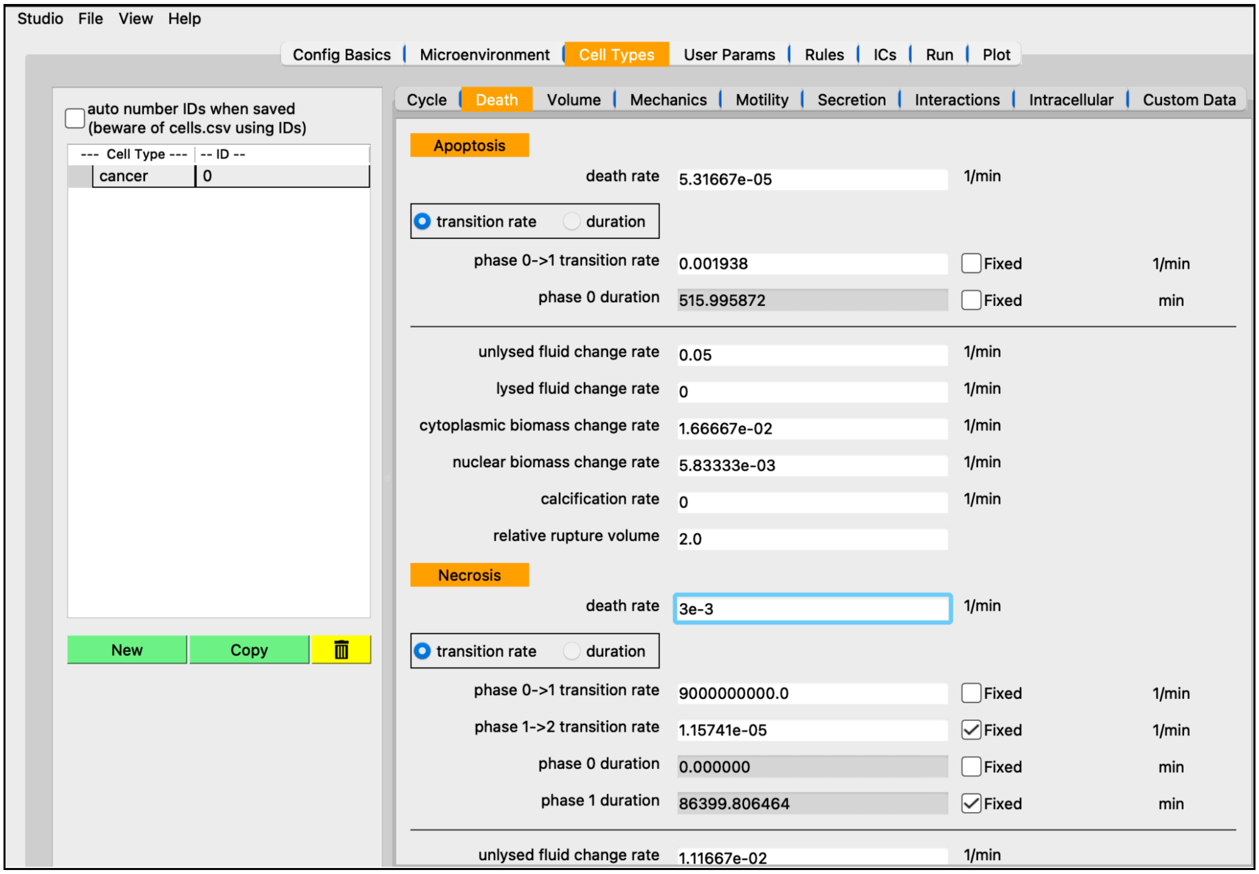


Figure 9. Editing Necrosis death rate.

- in the “Rules” tab, the “Cell Type” combobox should only contain “cancer”
- choose “oxygen” as the Signal
- chose “necrosis” as the Behavior
- choose “decreases” as the response
- choose 0.0 as the Saturation value of the Behavior
- choose 3.75 (mmHg) as the Half-max
- choose 8 as the Hill power
- optionally, Plot the new rule
- click “Add rule” and “Save”

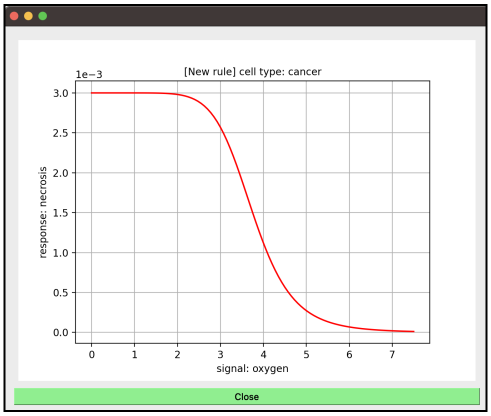


Figure 10. (oxygen, necrosis) Hill function.

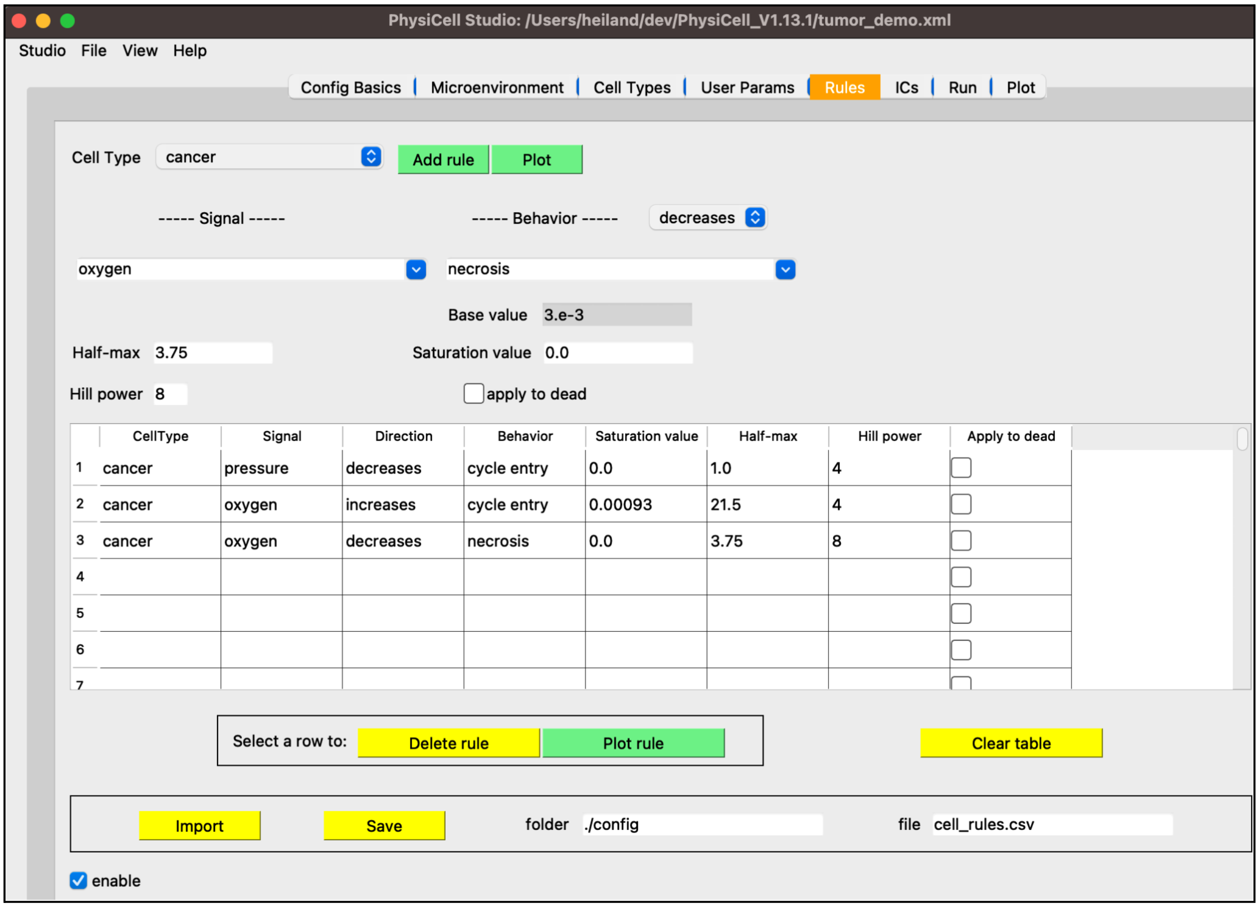


Figure 11. Rule #3: Oxygen decreases necrosis.

Add a more interesting boundary condition:

- in the “Microenvironment” tab:
    - set Dirichlet xmin=10 and ymin=10

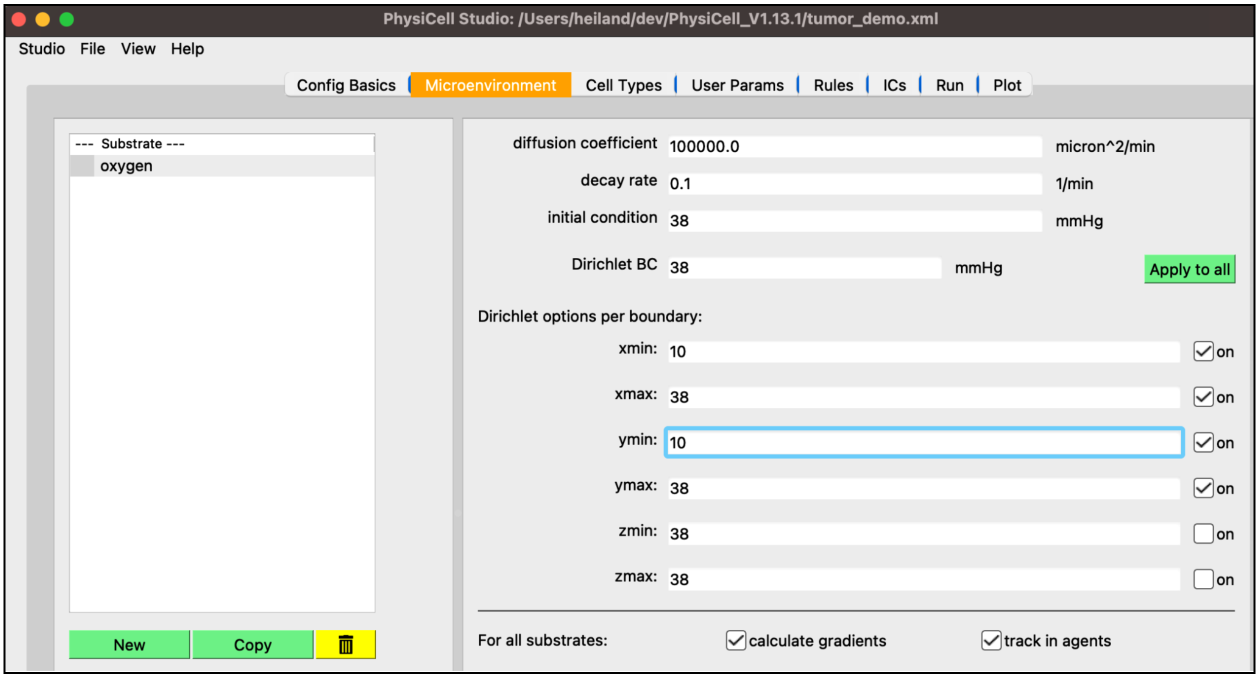

Figure 12. Editing Dirichlet conditions on xmin, ymin boundaries.

- Run a new simulation
- in the "Plot" tab, select ".svg" for cells (apoptotic cells are black, necrotic cells are brown)
- Play the simulation and notice:
  - cycling is preferential in the region with more oxygen
  - necrosis is preferential in the region with less oxygen

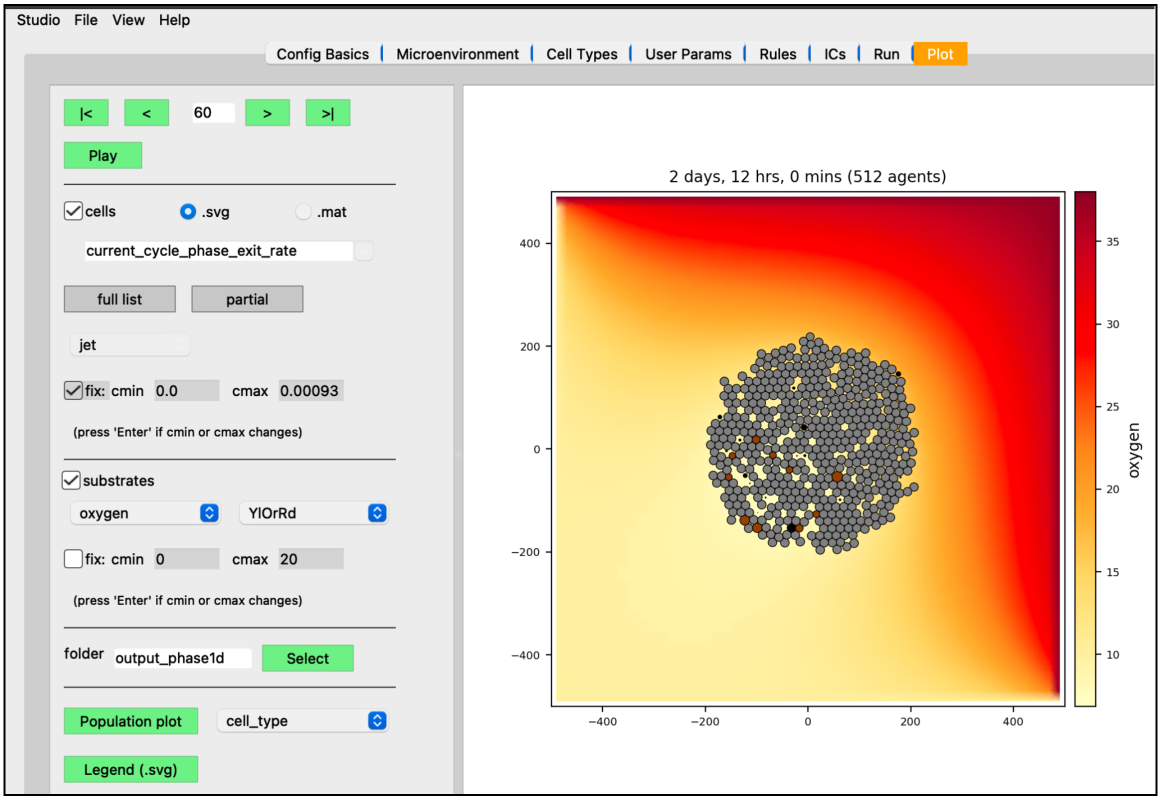


Figure 13. Results with three rules (plotting .svg cells and oxygen).

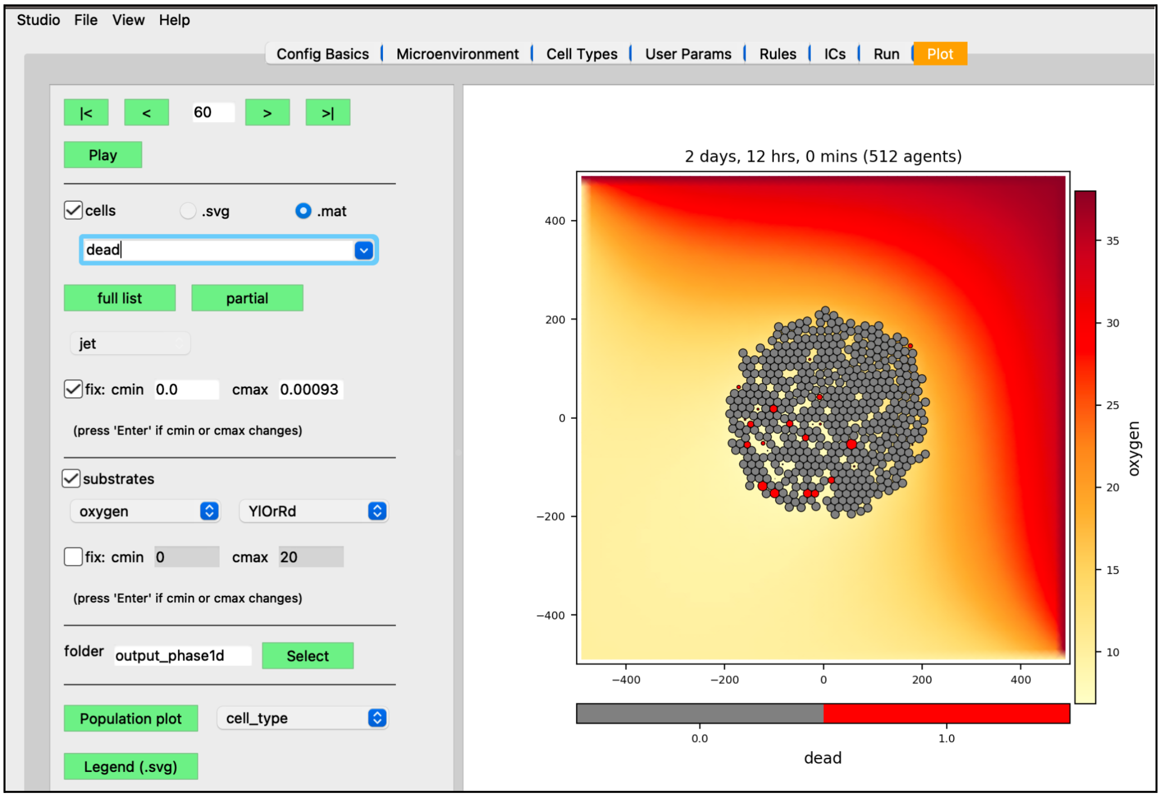


Figure 14. Alternatively, cells .mat option and color as dead (red) or alive (gray).

### Add a cytotoxic drug

- in the "Microenvironment" tab:
  - click "New", then select (e.g., double-click) the newly named one and rename it "drug" (and press Enter)
  - set its "diffusion coefficient" to 1600
  - set its "decay rate" to 0.002
  - set its "initial condition" to 10

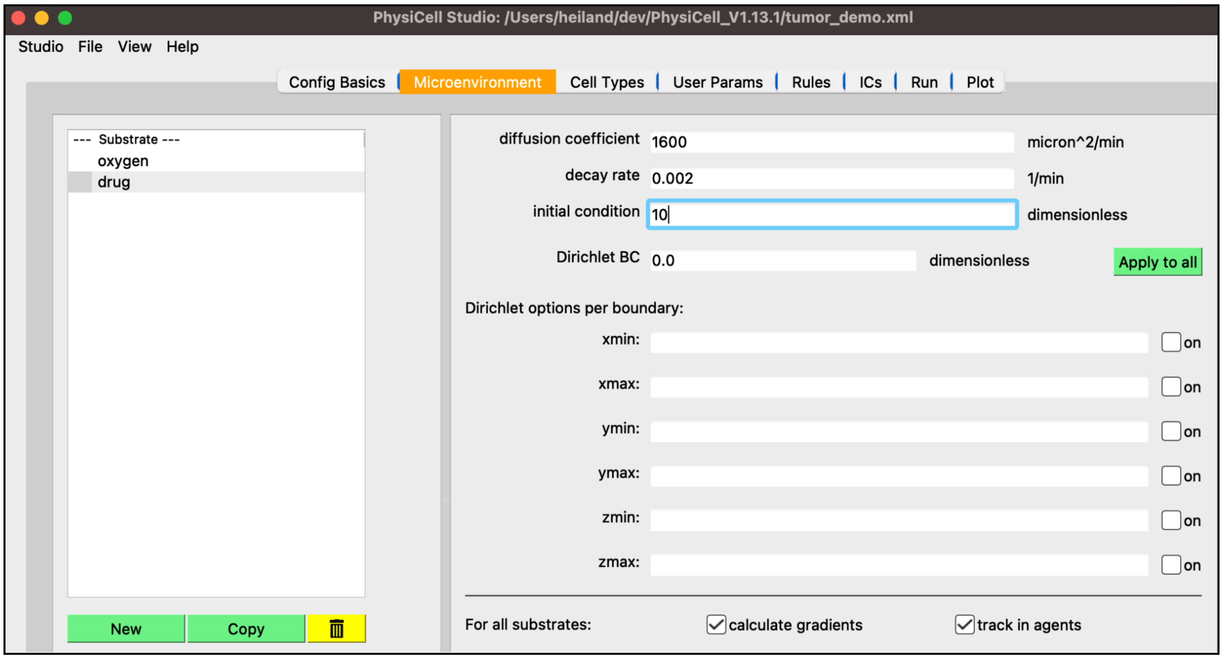


Figure 15. Edit drug parameters.

- in the "Cell Types" tab, "Secretion" sub-tab:
  - select "drug" in the combobox of substrates
  - set "uptake rate" to 1

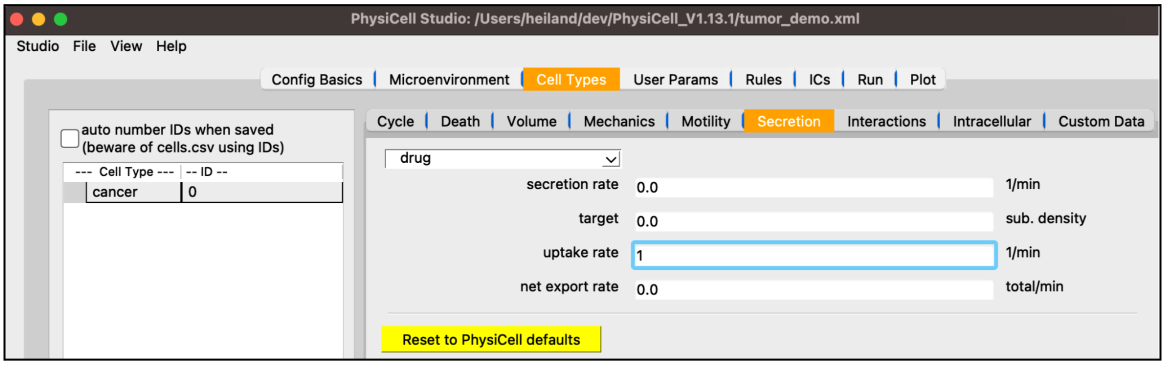


Figure 16. Edit drug uptake rate.

- in the “Rules” tab:
  - Cell Type: cancer
  - Signal: drug
  - Behavior: apoptosis  (response: increases)
  - Half-max: 0.5
  - Hill power: 4
  - Saturation value: 5e-3
  - Add rule: apoptosis increases with drug
  - Run simulation

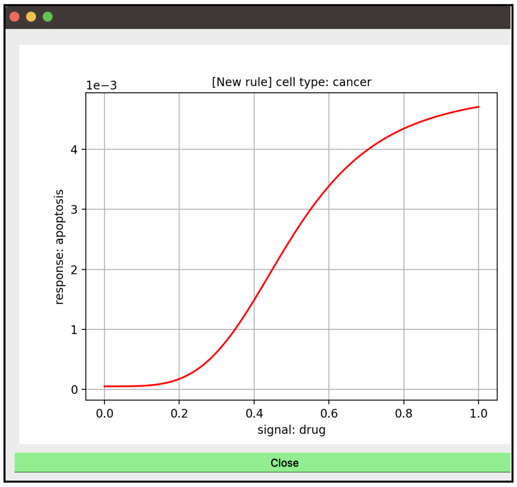


Figure 17. (drug, apoptosis) Hill function.

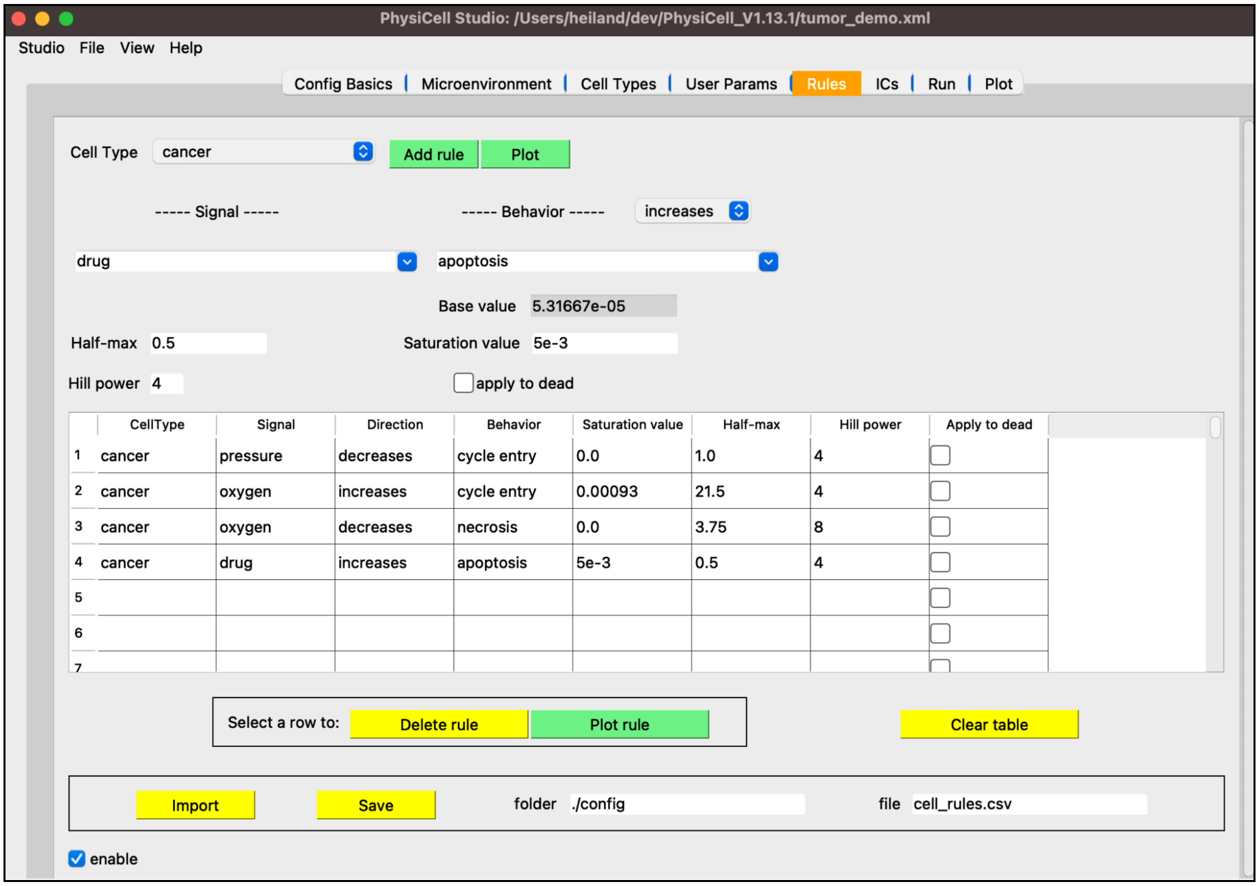


Figure 18. Rule #4: Drug increases apoptosis.

- in the "Plot" tab:
  - display cells as ".svg"
  - select "drug" substrate
  - Play to show results:
    - lots of cell death within first few hours
    - drug is quickly removed due to fast uptake by cells

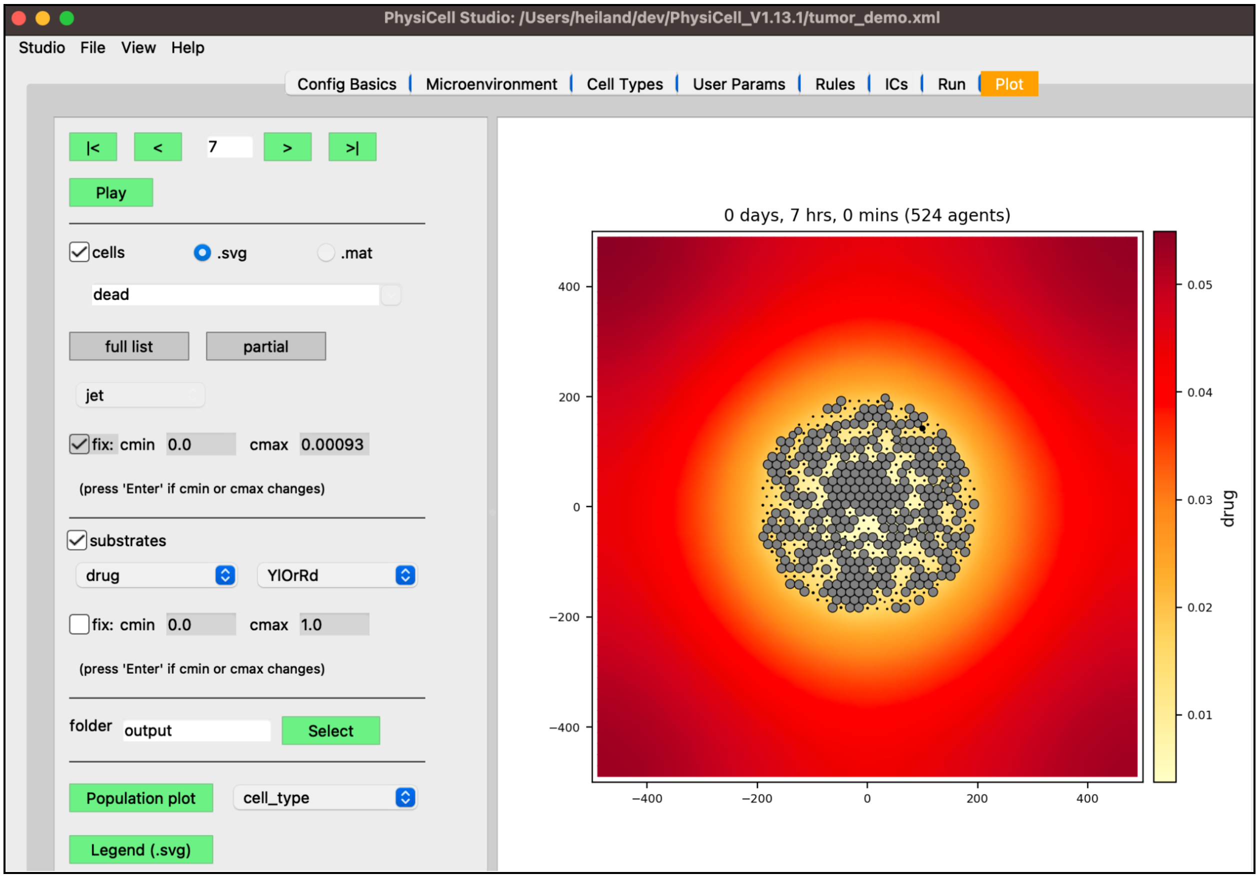


Figure 19. Results with four rules.

### Add release of dead cell debris

- in the "Microenvironment" tab:
  - click "New", then select (e.g., double-click) the newly named one and rename it "debris" (and press Enter)
  - set its "diffusion coefficient" to 1
  - set its "decay rate" to 0
  - set its "initial condition" to 0

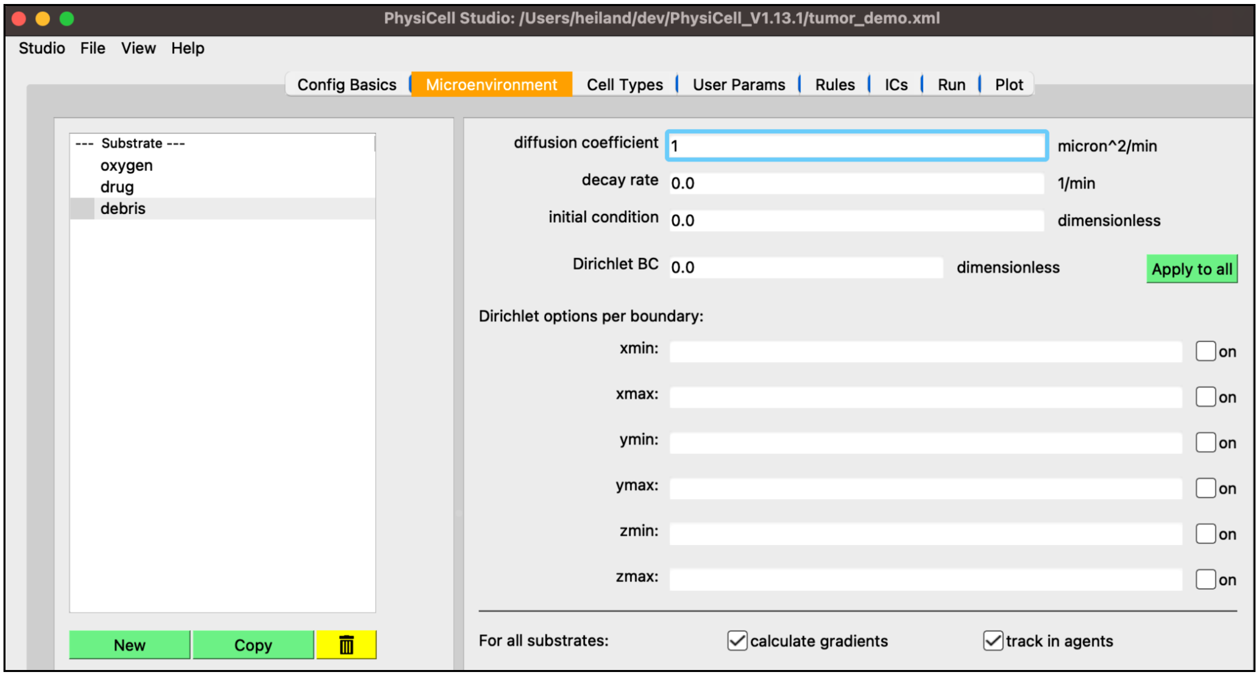


Figure 20. Edit debris parameters.

- in the "Cell Types" tab, "Secretion" sub-tab:
    - select "debris" in the combobox of substrates
    - set "target" to 1

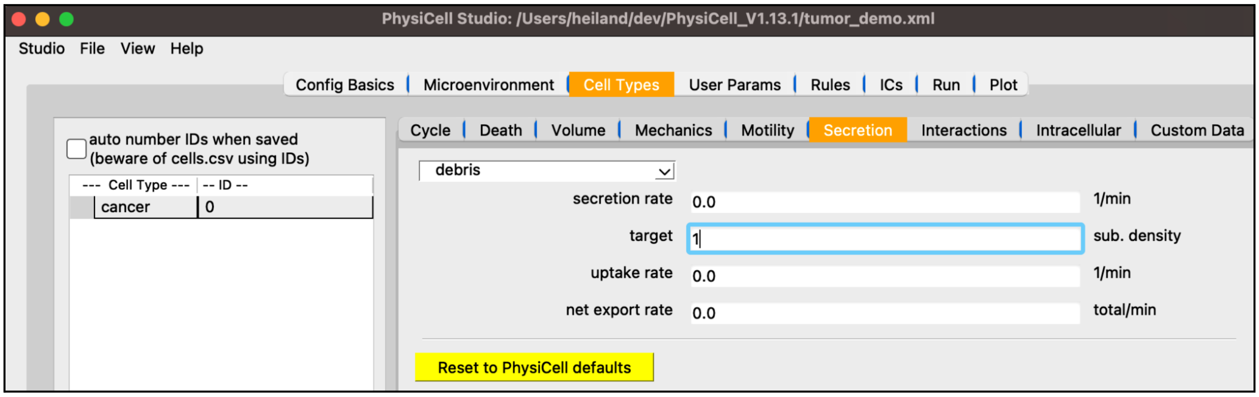


Figure 21. Edit target value for secretion of debris.

- in the "Rules" tab:
    - Cell Type: cancer
    - Signal: dead
    - Behavior: debris secretion (response: increases)
    - Half-max: 0.5
    - Hill power: 10

- Saturation value: 1
- apply to dead: true (check)
- Add rule: debris is secreted when a cell dies
- Run simulation

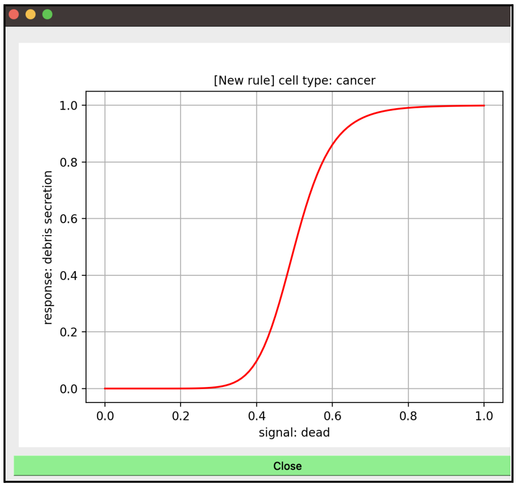


Figure 22. (dead, debris secretion) Hill function.

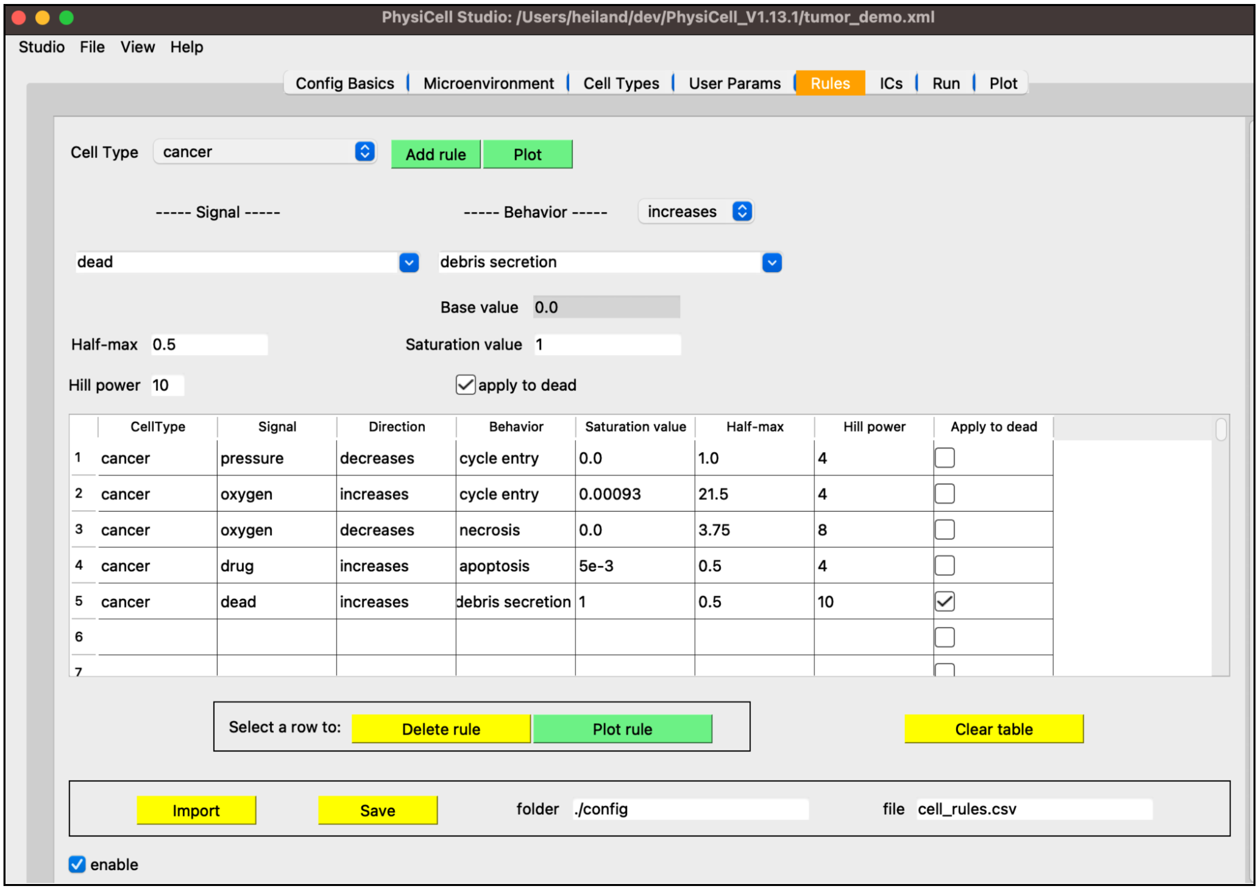


Figure 23. Death (dead signal) increases debris secretion.

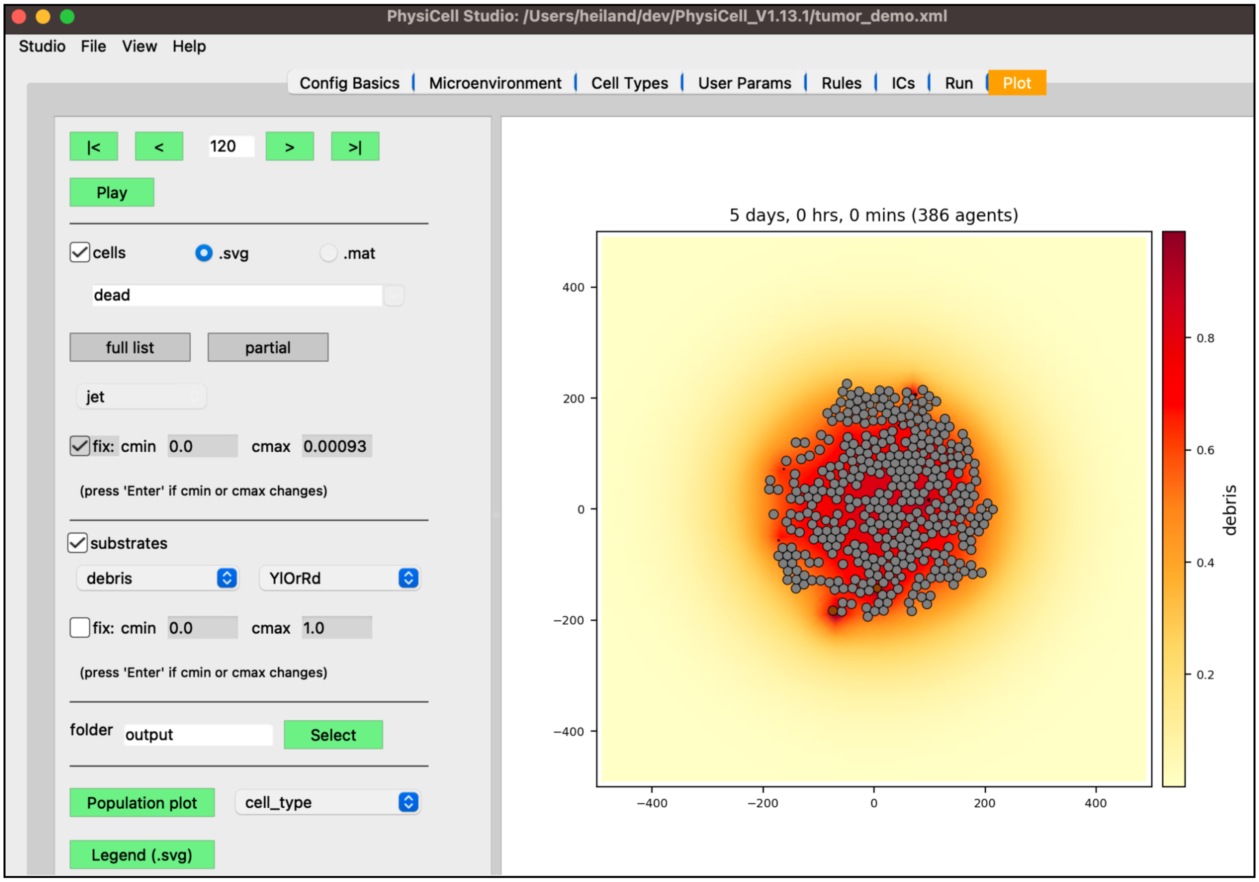


Figure 24. Results with five rules (.svg cells and debris signal).

Cell debris starts to accumulate in the region of the tumor.

### Add macrophages

- in the “Cell Types” tab:
  - click “cancer” cell type and click Copy
  - rename the copy to be “macrophage”
- in the “Death” sub-tab (with “macrophage” still selected):
  - set Apoptosis “death rate” to 0
  - set Necrosis “death rate” to 0

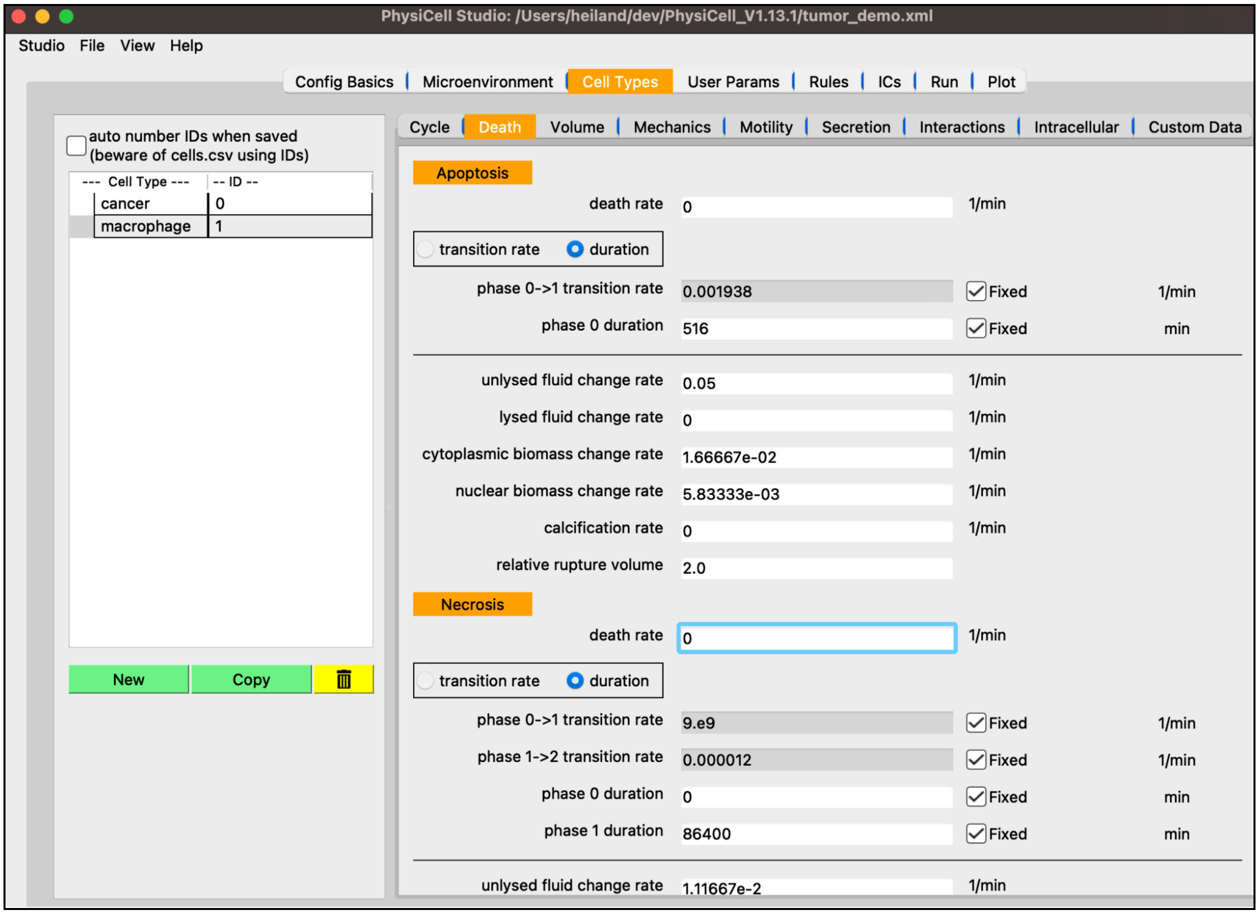


Figure 25. Create macrophage cell types with death rates=0.

- in the “Secretion” sub-tab (with “macrophage” still selected):
    - set oxygen “uptake rate” to 10
    - set drug “uptake rate” to 0
    - set debris “uptake rate” to 1

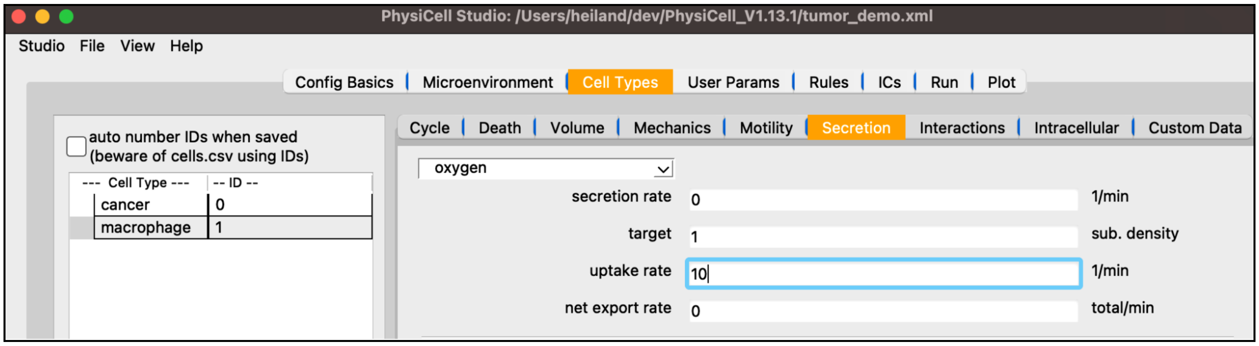


Figure 26. Edit macrophage secretion parameters for oxygen.

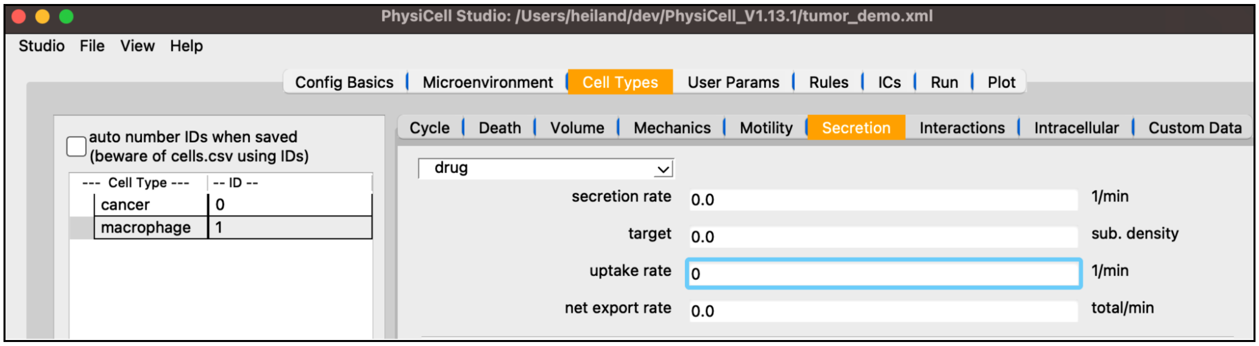


Figure 27. Edit macrophage secretion parameters for drug.

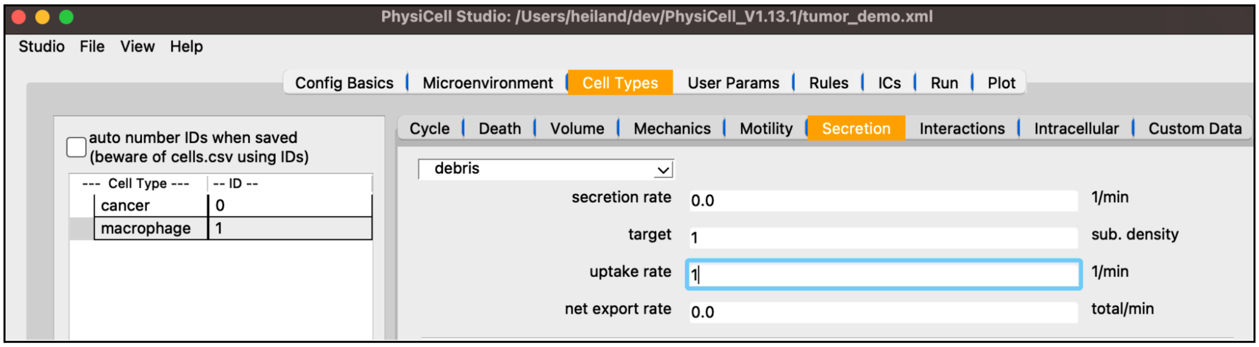


Figure 28. Edit macrophage secretion parameters for debris.

- in the "Mechanics" sub-tab (with "macrophage" still selected):
  - set "cell-cell adhesion strength" to 0
  - set "relative max adhesion distance" to 1.5

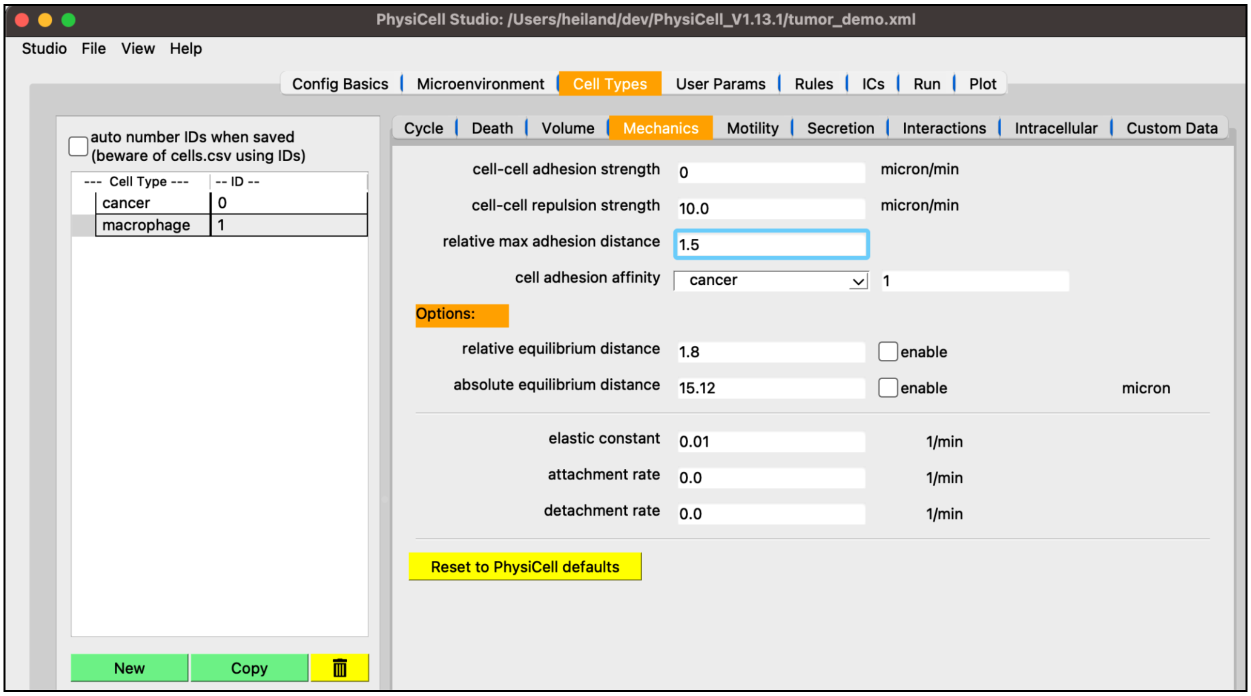

Figure 29. Edit macrophage mechanics parameters.

- in the “Motility” sub-tab (with “macrophage” still selected):
  - set “speed” to 1
  - set “persistence time” to 5
  - set “migration bias” to 0.5
  - check “enable motility”
  - Chemotaxis: check “enabled” and “towards” debris

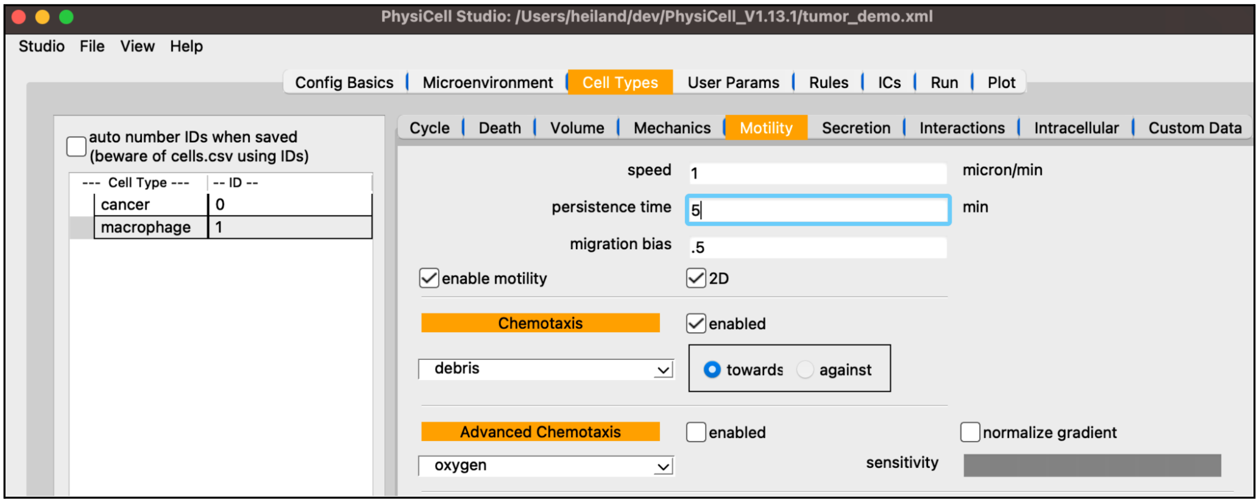

Figure 30. Edit macrophage motility parameters.

- in the “Interactions” sub-tab (with “macrophage” still selected):
    - set “dead phagocytosis rate” to 0.01

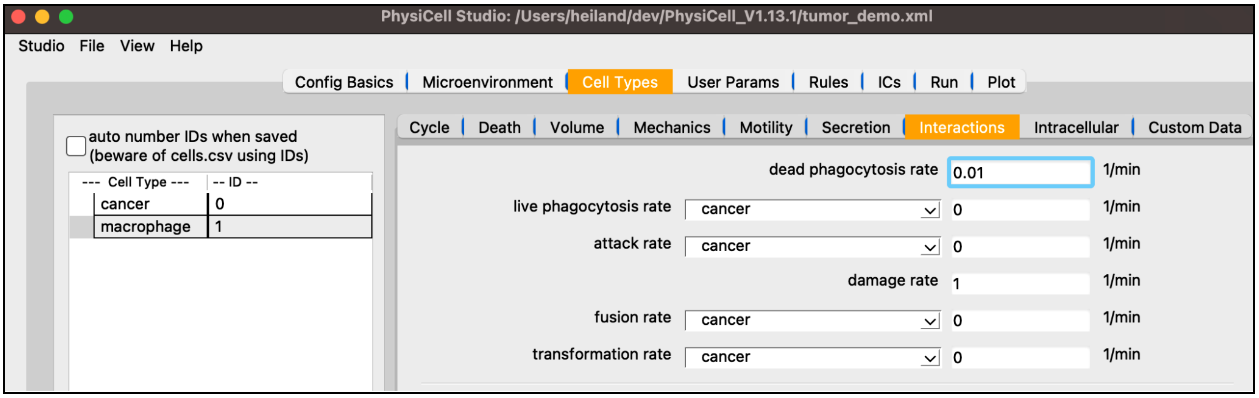


Figure 31. Edit macrophage phagocytosis rate.

- in the “ICs” tab:
    - select “cell type” to be macrophage
    - select “annulus/disk”
    - select “random fill”
    - # cells = 100
    - set R1 (min radius) = 250
    - set R2 (max radius) = 300
    - click Plot (creates an annulus of randomly placed macrophage cells)
    - click Save

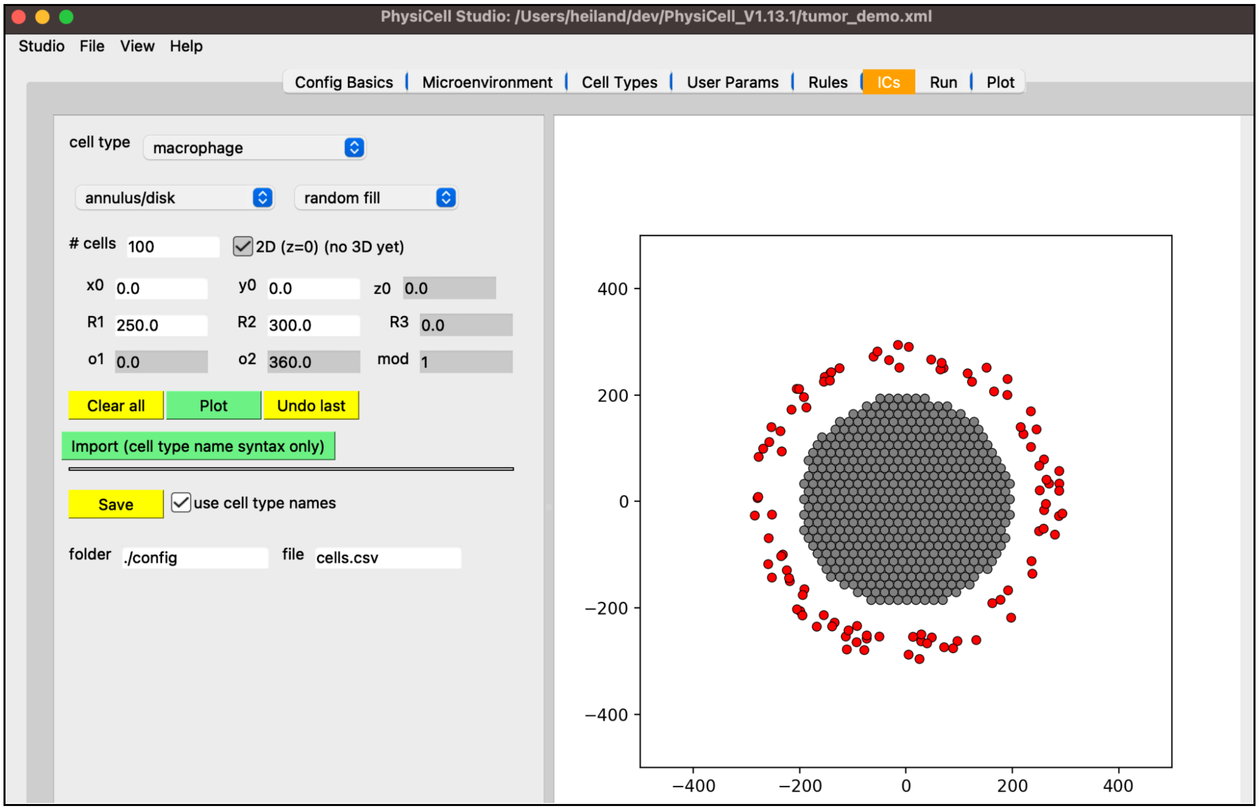


Figure 32. Generate macrophage cells' initial positions.

- Run a simulation (note we did not create any rule using macrophages yet)
- in the Plot tab:
    - display cells as ".svg"
    - display "debris" substrate; select "viridis" colormap
    - View menu → Plot options: uncheck "fill" for Cells (to see the debris better)

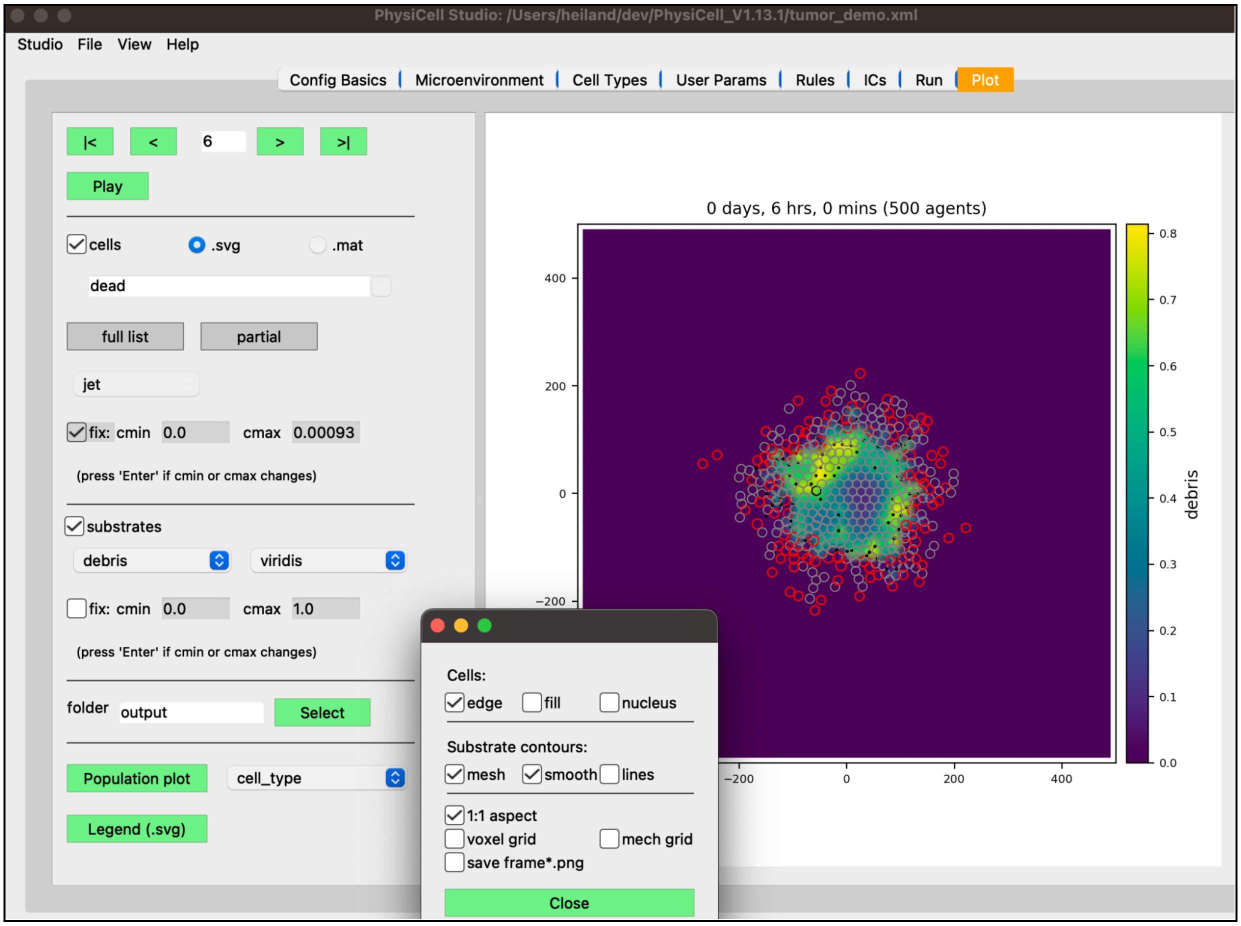


Figure 33. Debris field at 6 hours (uncheck filled cells to see better).

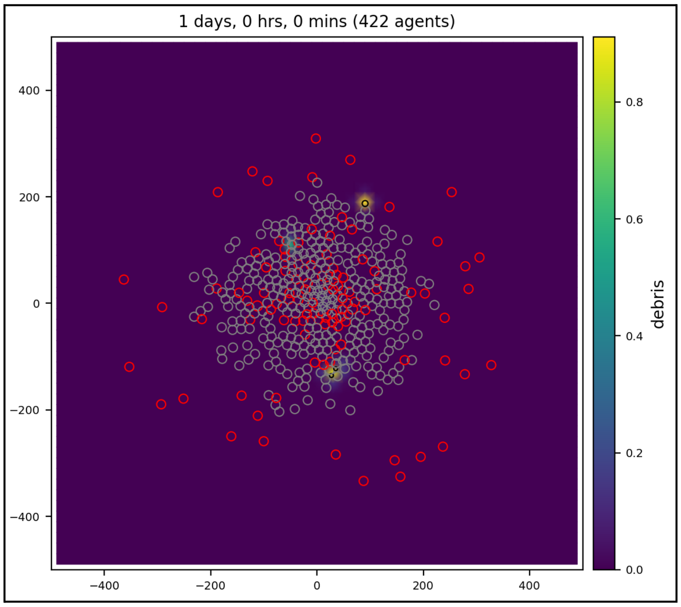


Figure 34. Debris field at 24 hours.

### Add pro-inflammatory factor

- in the “Microenvironment” tab:
    - click “New”, then select (e.g., double-click) the newly named one and rename it “pro-inflammatory factor” (and press Enter)
    - set its “diffusion coefficient” to 1000
    - set its “decay rate” to 0.1
    - set its “initial condition” to 0

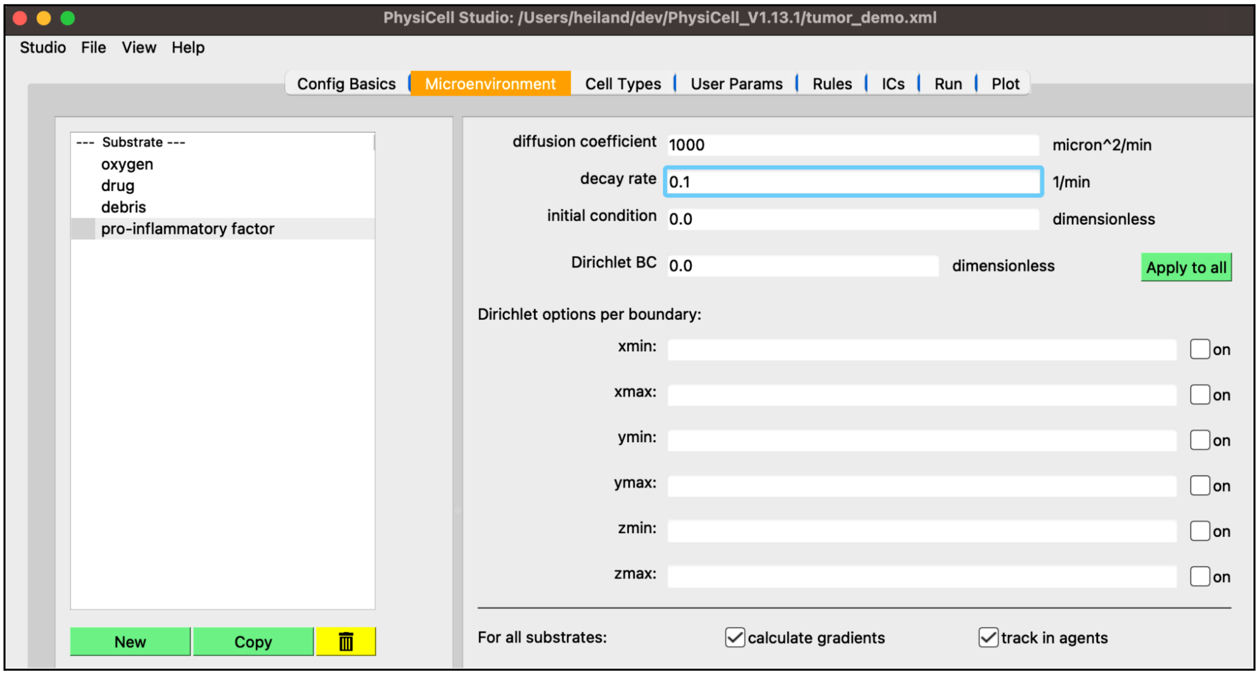


Figure 35. Create pro-inflammatory factor signal and set its parameters.

- in the “Cell Types” tab:
    - click “macrophage” cell type and click Copy
    - rename the copy to be “M1 macrophage”
- in the “Secretion” sub-tab (with “M1 macrophage” still selected):
    - select “pro-inflammatory factor” as the substrate
    - set “secretion rate” to 10
    - set “target” to 1

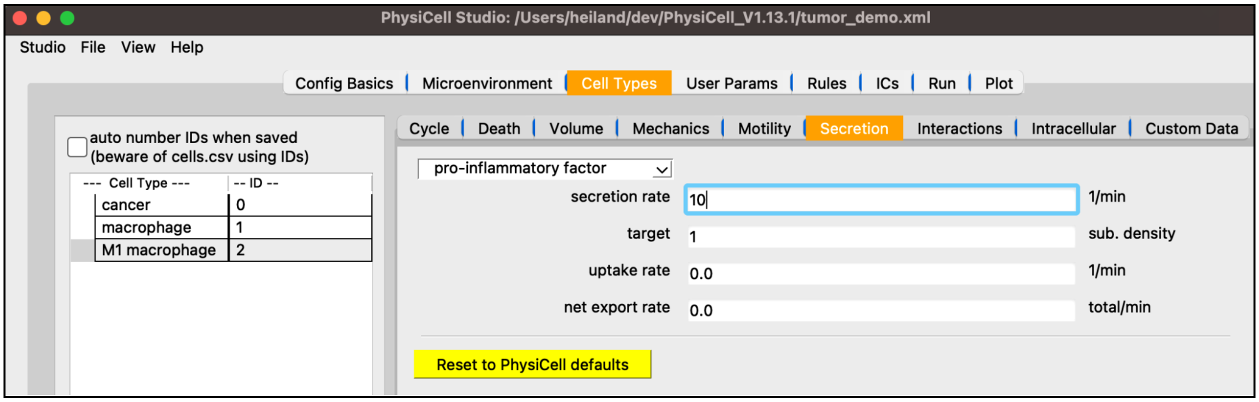


Figure 36. Create M1 macrophage cell type and set its secretion parameters.

- in the “Rules” tab:
  - Cell Type: macrophage
  - Signal: contact with dead cell
  - Behavior: transform to M1 macrophage (response: increases)
  - Half-max: 0.5
  - Hill power: 10
  - Saturation value: 0.1
  - apply to dead: false (not checked)
  - Add rule: macrophage is transformed to M1 macrophage upon contact with dead cell
  - Run simulation

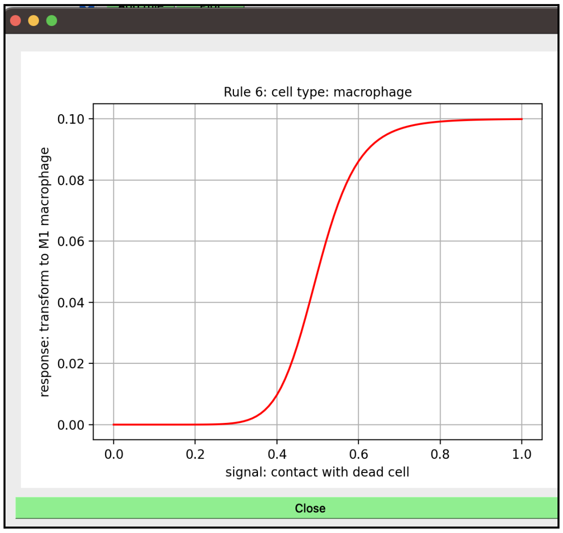


Figure 37. (contact with dead cell, transform to M1 macrophage) Hill function (for macrophage).

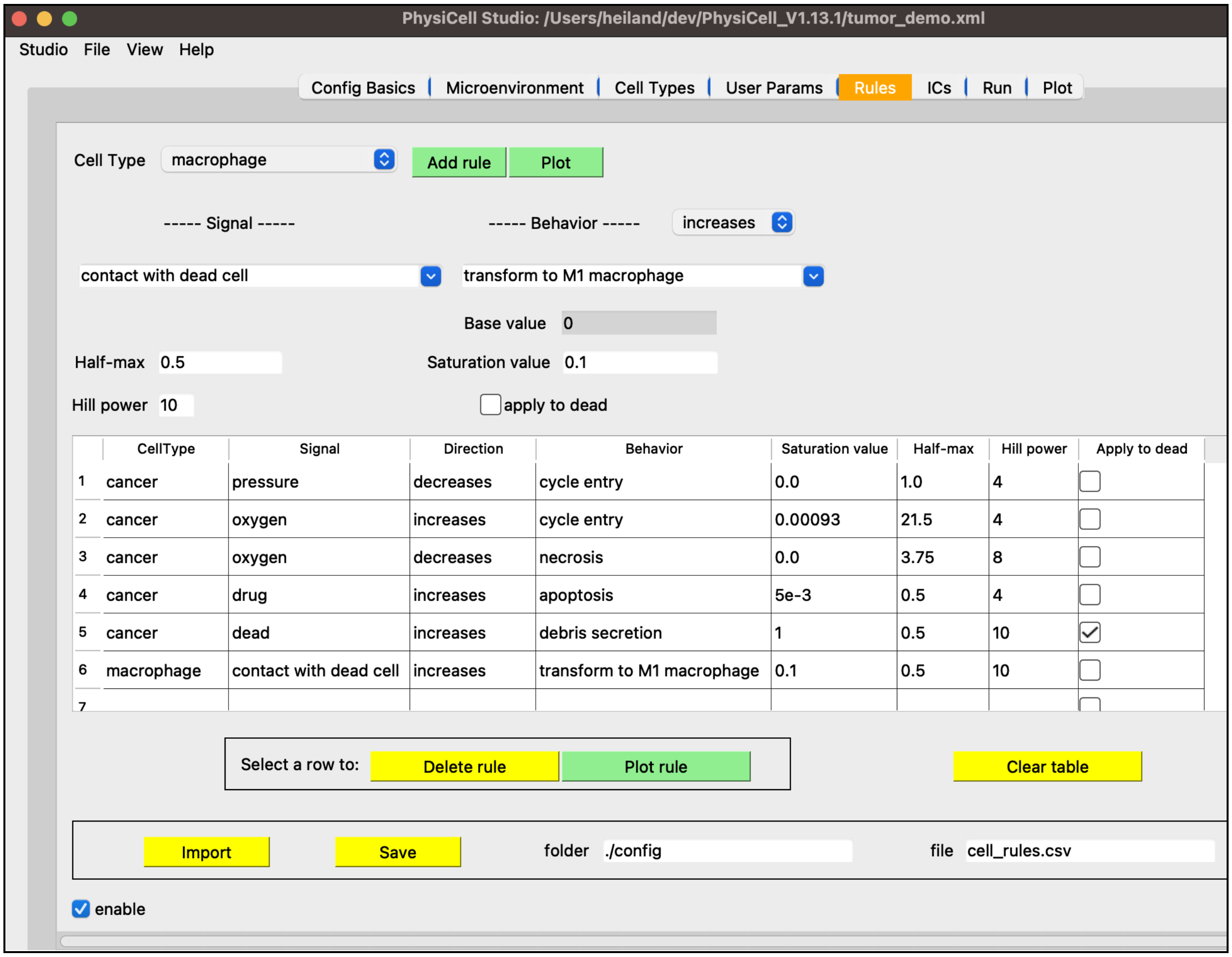


Figure 38. Rule #6: Macrophage contact with dead cell increases transform to M1 macrophage.

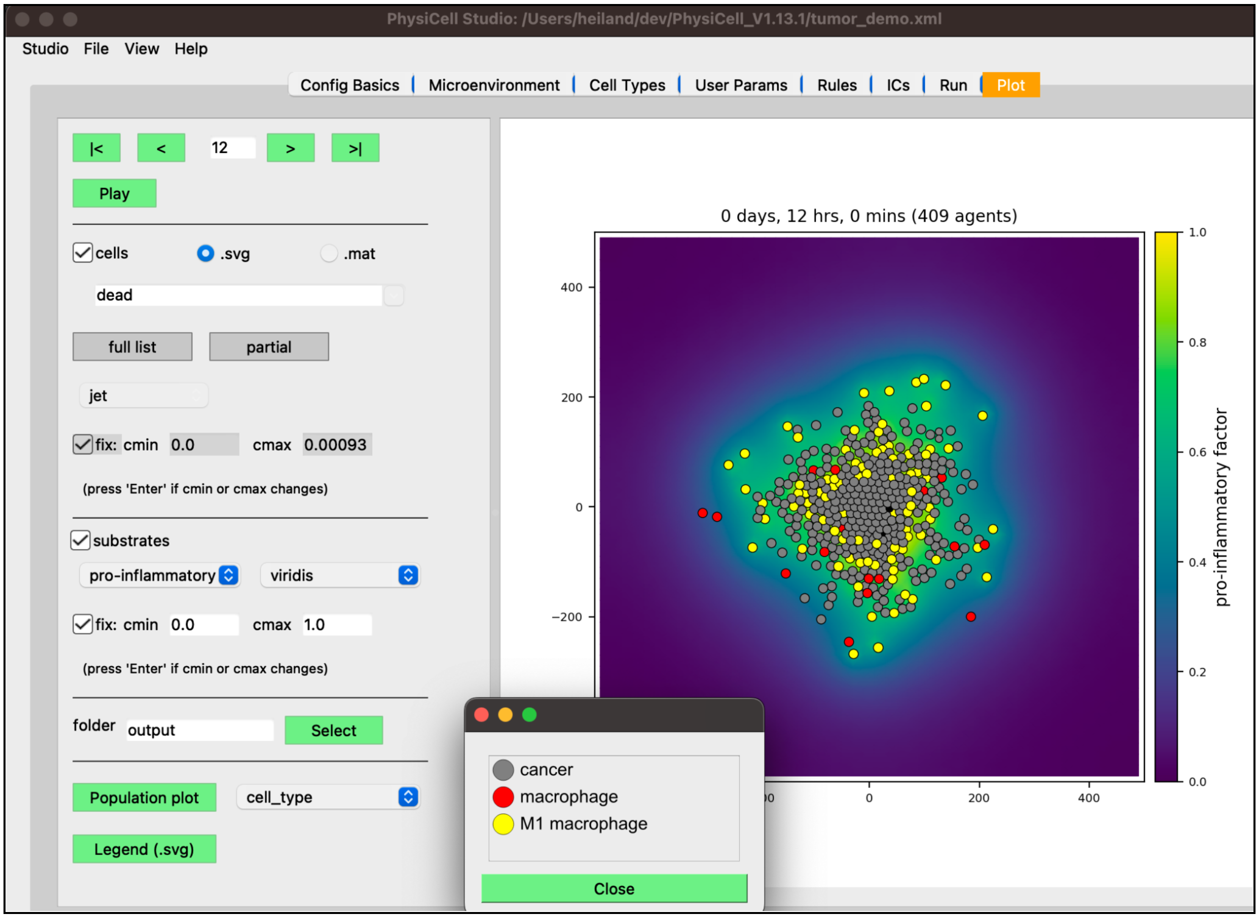


Figure 39. Results using six rules, at 12 hours.

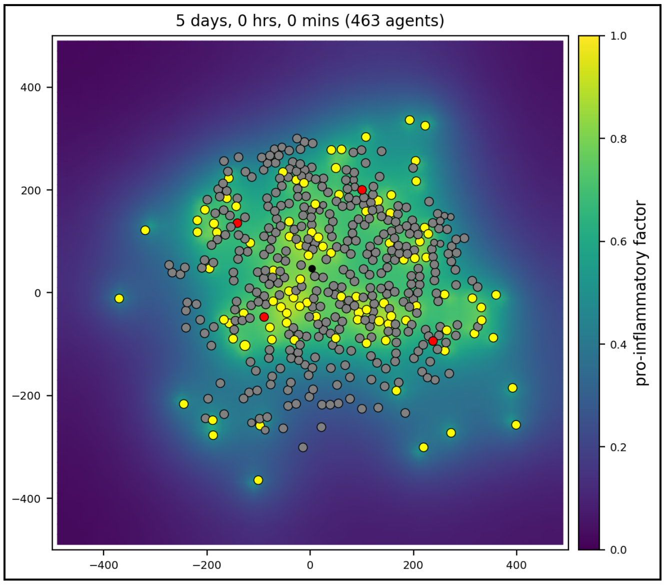


Figure 40. Results at 5 days.

### Add effector T cells

- in the “Cell Types” tab:
  - click “macrophage” cell type and click Copy
  - rename the copy to be “Effector T cell”
- in the “Motility” sub-tab (with “Effector T cell” still selected):
  - select “pro-inflammatory factor” as the substrate in Chemotaxis

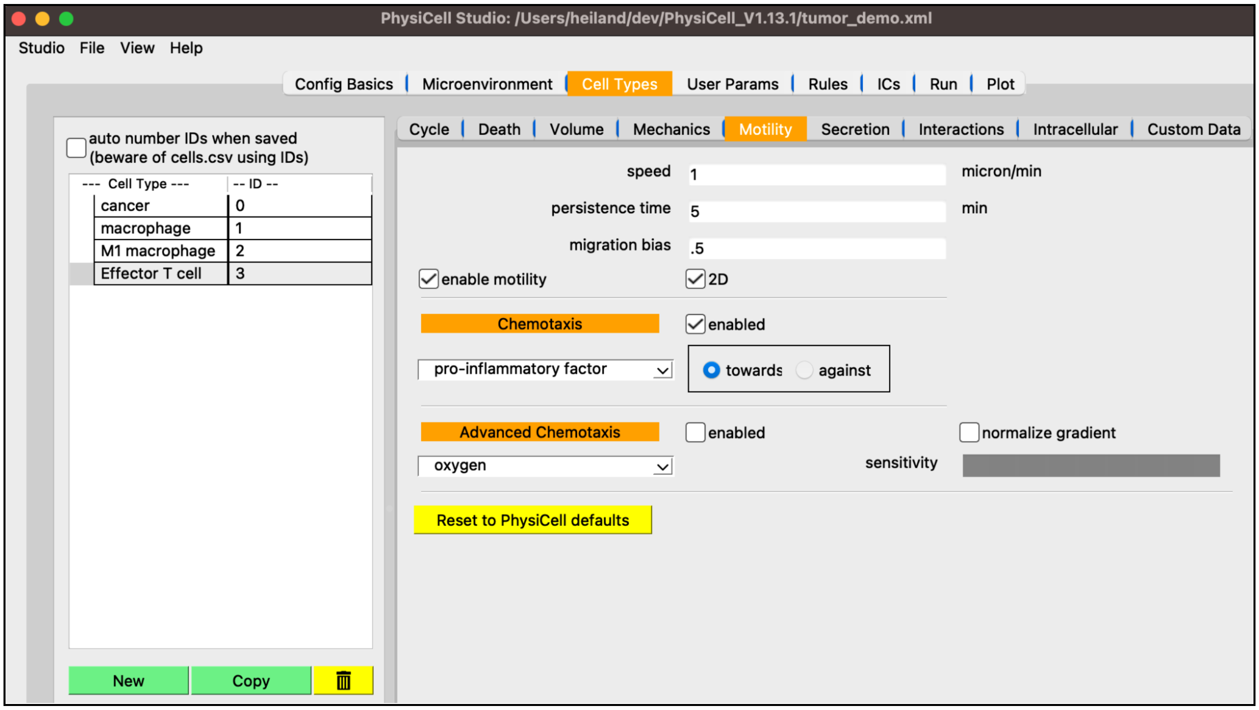


Figure 41. Create “Effector T cell” cell type.

- in the “Secretion” sub-tab (with “Effector T cell” still selected):
  - select “debris” as the substrate
  - set “uptake rate” to 0
  - select “pro-inflammatory factor” as the substrate
  - set “uptake rate” to 1

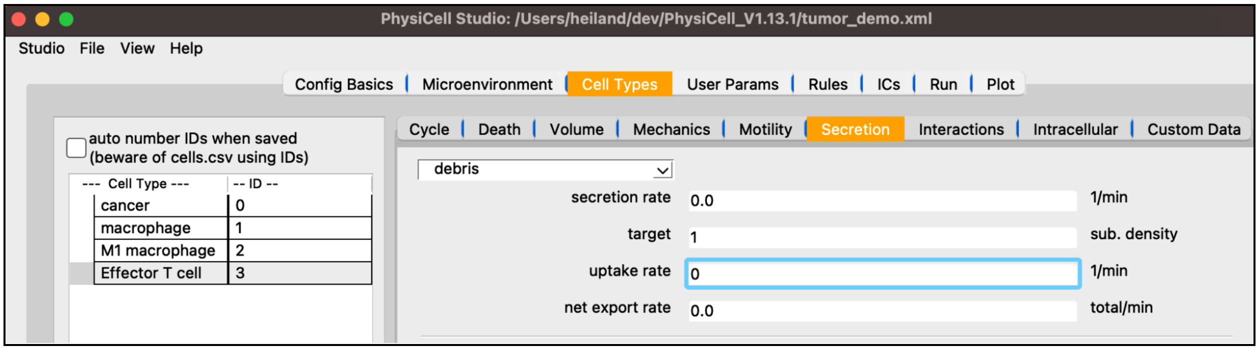


Figure 42. Edit “Effector T cell” secretion parameters for debris.

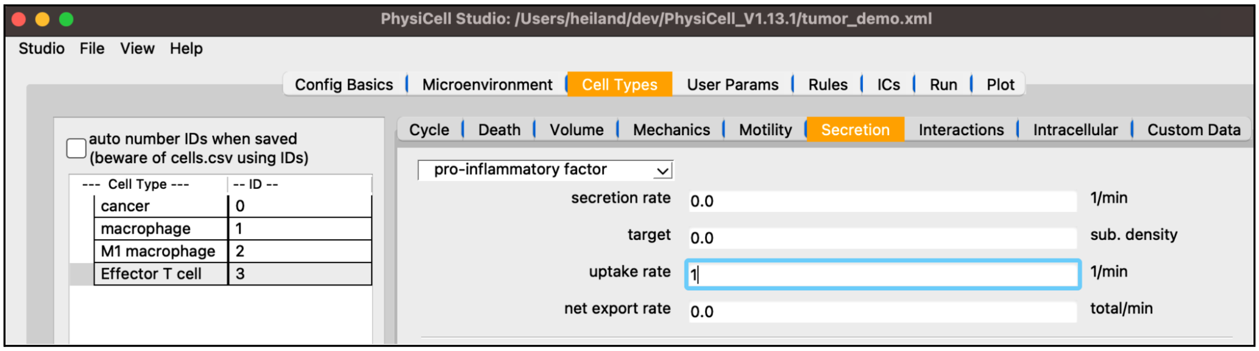


Figure 43. Edit “Effector T cell” secretion parameters for pro-inflammatory factor.

- in the “Interactions” sub-tab (with “Effector T cell” still selected):
  - set “dead phagocytosis rate” to 0
  - set “attack rate” for cancer to 10

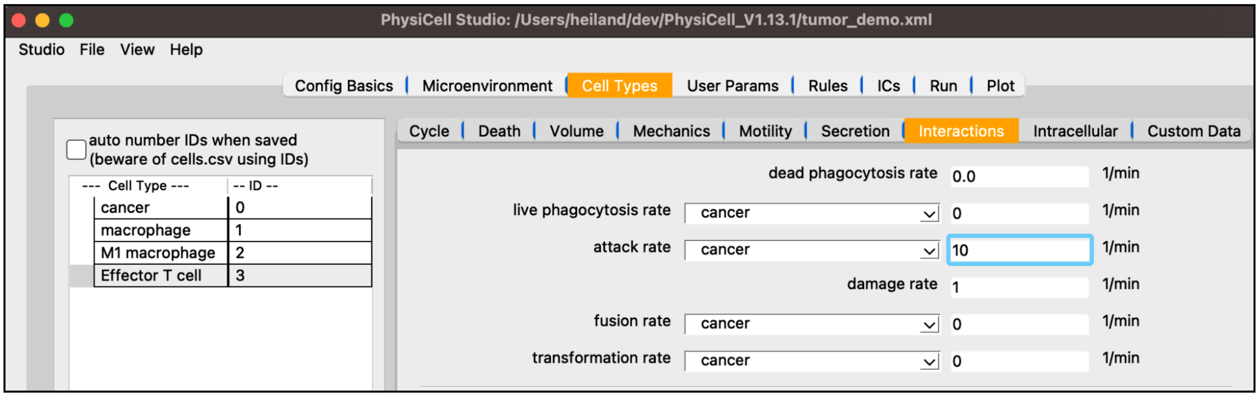


Figure 44. Edit “Effector T cell” Interaction parameters.

- in the “Rules” tab:
  - Cell Type: cancer
  - Signal: damage
  - Behavior: apoptosis (response: increases)

- Half-max: 5
- Hill power: 4
- Saturation value: 0.01
- apply to dead: false (not checked)
- Add rule: cancer cell damage increases apoptosis

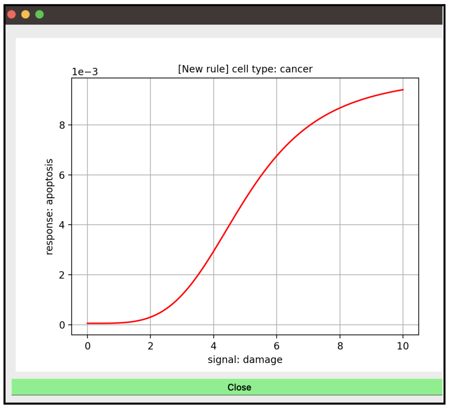


Figure 45. (damage, apoptosis) Hill function (for cancer cell).

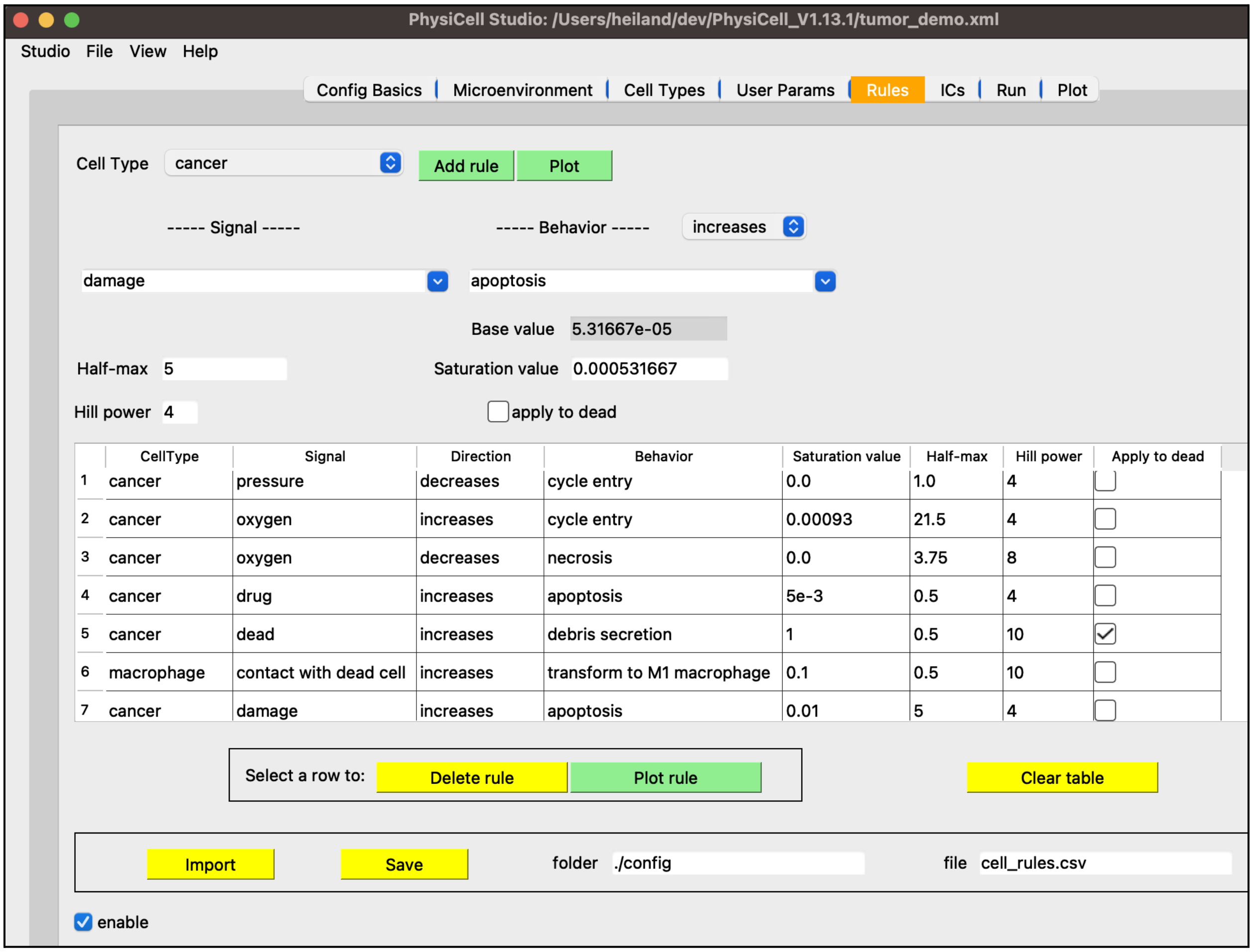


Figure 46. Rule #7: damage increases apoptosis (of cancer cell).

- in the “ICs” tab:
    - select “cell type” to be Effector T cell
    - select “annulus/disk”
    - select “random fill”
    - # cells = 100
    - set R1 (min radius) = 250
    - set R2 (max radius) = 300
    - click Plot
    - click Save

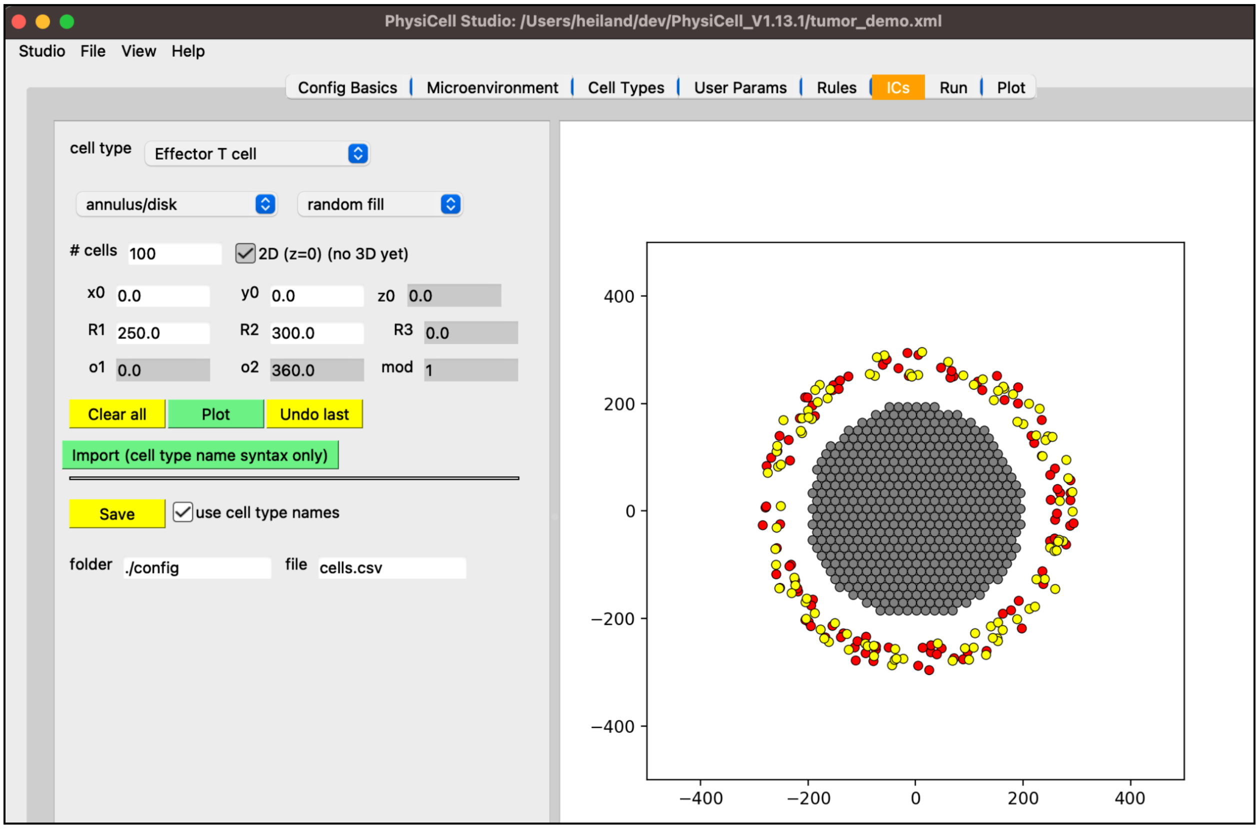


Figure 47. Generate Effector T cells initial positions.

- Run simulation
- in the Plot tab:
    - select “.svg” for cells (remember, apoptotic cells are black, necrotic are brown)
    - Play the simulation
    - note how Effector T cells attack and kill off cancer cells

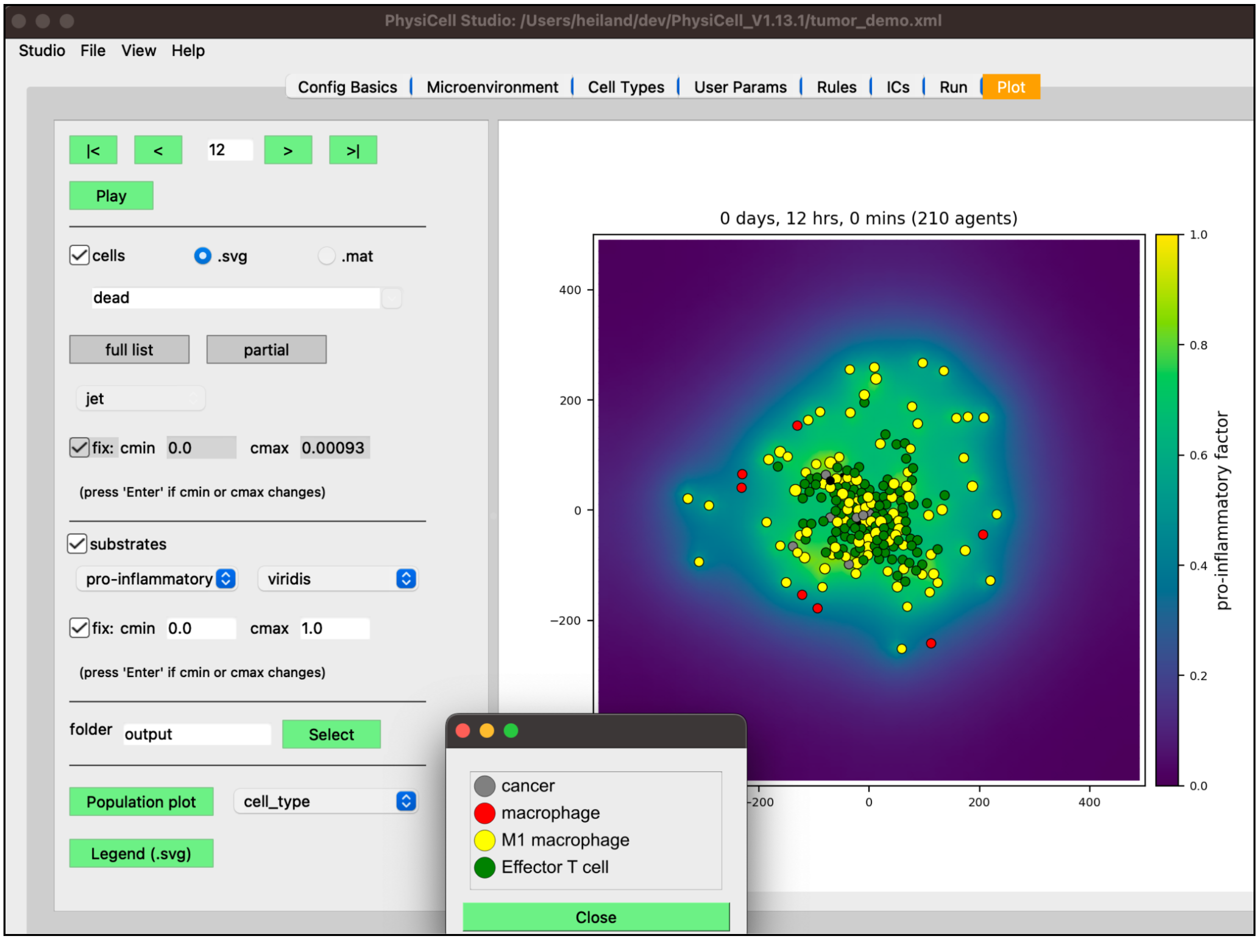


Figure 48. Results using seven rules, at 12 hours, showing pro-inflammatory factor (with fixed colormap range.

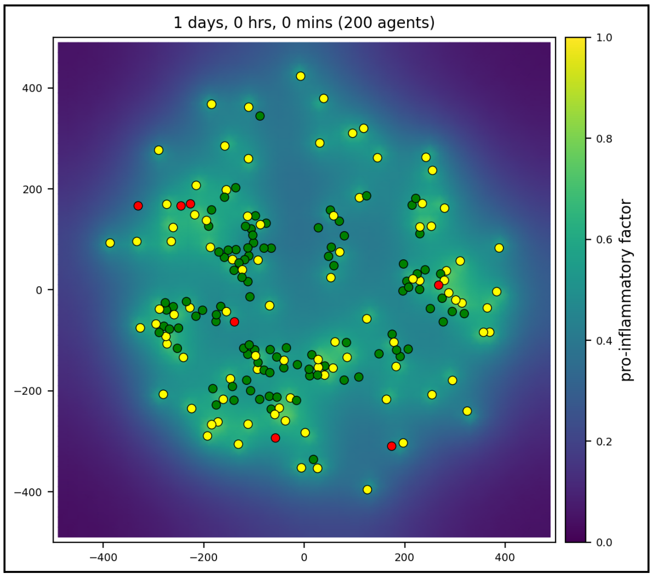


Figure 49. Results at 24 hours.

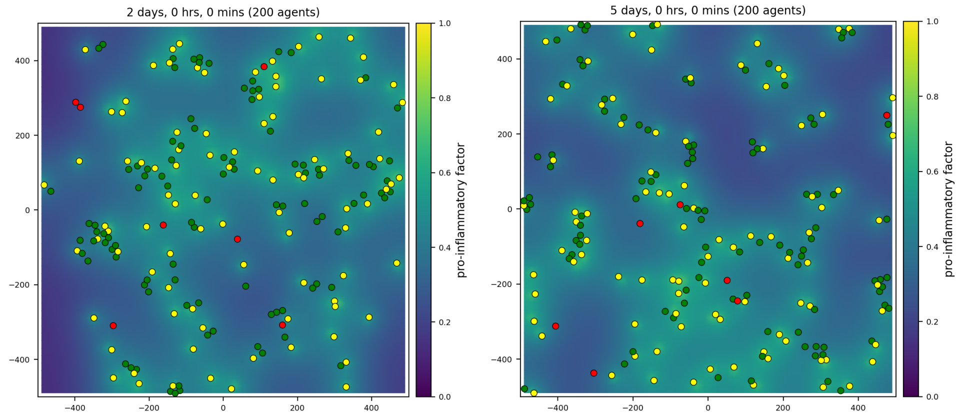


Figure 50. Results at 2 days (left) and 5 days (right).

### Intracellular modeling

PhysiCell Studio also provides interfaces to intracellular modeling. Although intracellular modeling is not used in the tumor model presented above, we wanted to present it in this Supplemental material. Currently, only a boolean intracellular modeling[2] interface is provided in the Studio. In the future, we will also provide an interface for ODE intracellular models[3].

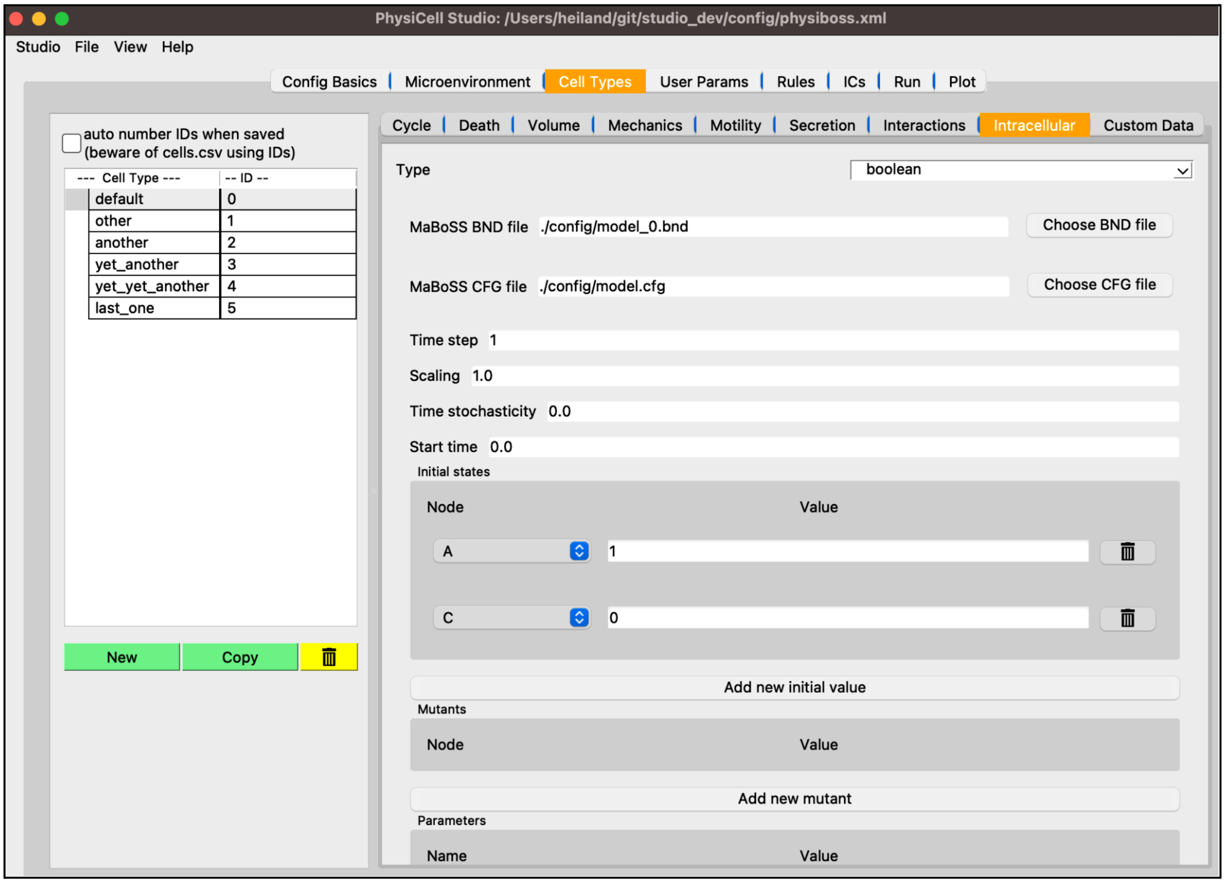


Figure 51. Interface for a boolean intracellular model.

1. Jeanette A.I. Johnson, Genevieve L Stein-O'Brien, Max Booth, Randy Heiland, Furkan Kurtoglu, Daniel Bergman, et al. Digitize your Biology! Modeling multicellular systems through interpretable cell behavior. bioRxiv. 2023; 2023.09.17.557982.

2. Ponce-de-Leon M, Montagud A, Noël V, Pradas G, Meert A, Barillot E, et al. PhysiBoSS 2.0: a sustainable integration of stochastic Boolean and agent-based modelling frameworks. bioRxiv. 2023. p. 2022.01.06.468363. doi:10.1101/2022.01.06.468363

3. Somogyi ET, Bouteiller J-M, Glazier JA, König M, Medley K, Swat MH, et al. libRoadRunner: A High Performance SBML Simulation and Analysis Library. arXiv [q-bio.SC]. 2015. Available: http://arxiv.org/abs/1503.01095
